# Pro-drug peptide and its metabolites disrupt amyloid fibrils by destabilizing salt bridge interaction and planar beta-sheet topology

**DOI:** 10.1101/2020.09.09.290643

**Authors:** Vasista Adupa, Bhubaneswar Mandal, K. Anki Reddy

## Abstract

The most common age-related neurodegenerative disorder, Alzheimer’s disease, is clinically characterized by continuous neuronal loss resulting in loss of memory and dementia with no cure to date. Amyloid-*β* (A*β*) aggregates and tau protein are believed to be the causative agents of this pathogenesis. In the present study, we have investigated the effect of the Pro-Drug peptide (PDp) and its metabolites (*α*-aspartyl & *β*-aspartyl) on the A*β* aggregates using atomistic molecular dynamics simulations. One of the key findings in our work is in the presence of *α*-aspartyl as a ligand, the salt bridges which hold the N-terminals together are completely disrupted, thus setting the N-terminals free and exposed entirely to the solvent which can make the aggregation of A*β* less severe. The efficiency of the ligands, which are responsible for the disruption of A*β*, depends on the alignment and strength of the repulsive interactions. Besides repulsive interactions, we found that there is a need for hydrogen bonding, which acts as a support for the ligand to stay in the vicinity of the aggregate. Moreover, we have noticed that one of the metabolites, namely *β*-aspartyl, formed more hydrogen bonds with the aggregate than the other ligands and had a different mode of action with the chains of A*β* due to its unique flexible kink in the backbone.

## 0.1 Introduction

Advances in therapeutic design for Alzheimer’s disease (AD) include using synthetic peptides that inhibit fibrillogenesis of amyloid *β*-protein^*1*^ (A*β*), which is a seminal pathogenetic event in AD. One of the important strategies in designing these peptides was developed by Ghanta *et al.^2^* in which a recognition element which specifically binds to a part of A*β* and a structure-disrupting element that alters the A*β* aggregation pathways are added. Hilbich *et al.^3^* demonstrated that the hydrophobic residues (Leu^17^ to Phe^20^) are crucial for the fibril formation and structural alterations in this region may play a pivotal role in the cure of AD. Therefore peptides were developed by incorporating a short A*β* fragment KLVFF (A*β*^16–20^) that could bind to full-length A*β* and further prevent it from forming fibrils.^*4*^ Some of the other strategies explored in targeting A*β* are using antibodies^*5*^ and small molecules^*6–13*^.

Soto *et al.^14^* introduced *β*-Sheet Breaker peptides (BSBp) which contains a stretch of five amino-acids, where LVFF (A*β*^17-20^) was used as a template for designing the BSBp (LPFFD), where proline (P) is used as a structure disrupting element. Viet *et al.^15^* studied the effect of KLVFF and LPFFD on the oligomerization of amyloid peptides by all atoms molecular simulations and found out that LPFFD has more substantial influence than KLVFF on the aggregation of A*β*^16-22^. A previous MD study by bruce *et al.* found that the peptide LHFFD had a weaker interaction with the fibril than the peptide LPFFD.

In the present study, Pro Drug peptide (PDp), developed by Nadimpally *et al.^16^* and its metabolites are used as the ligands. The sequence of PDp is Ac-LD(OBzl)FFD-NH_2_, and it bears some unique features. PDp carries a sequence homology with A*β*^19-20^ (-F-F-) but is initially free from structure disrupting element. Thus, it is better for recognizing A*β* peptide than usual BSBp’s that bear sequence homology as well as the disrupting element inbuilt. However, in physiological pH and temperature, O to N acyl migration in the dipeptide (-D(OBzl)-F-) of PDp takes place, leading to the formation of aspartimide derivative, which further hydrolyse into *α* & *β*-aspartyl derivatives. These aspartimide derivatives, as well as the *α* & *β*-aspartyl derivatives, contains structure destabilizing elements and also bears sequence homology with A*β*. Thus, the in situ generated metabolites now behave as the mixture of the usual BSBp’s. These PDp’s are known to disrupt existing A*β* fibril^*17*^ and ameliorate A*β* protofibril mediated cytotoxicity in both cell based assay^*18*^ and animal models.^*19*^

Simulating the effect of such a ligand that changes its chemical structure with time (Scheme 1) is quite cumbersome using classical molecular dynamics. Therefore, we came up with an approach that can be well assumed as simulating a dynamic system. For this, we simulated the effect of PDp; it’s *α*-aspartyl derivative, *β*-aspartyl derivative, and their mixture, individually in parallel, such that the results of all the individual simulations collectively can be considered as simulating the dynamic system. Limitations of docking^*20*^ are considered, and the initial setup of the simulation was created by placing ligands randomly around the protein.

**Scheme 1:**
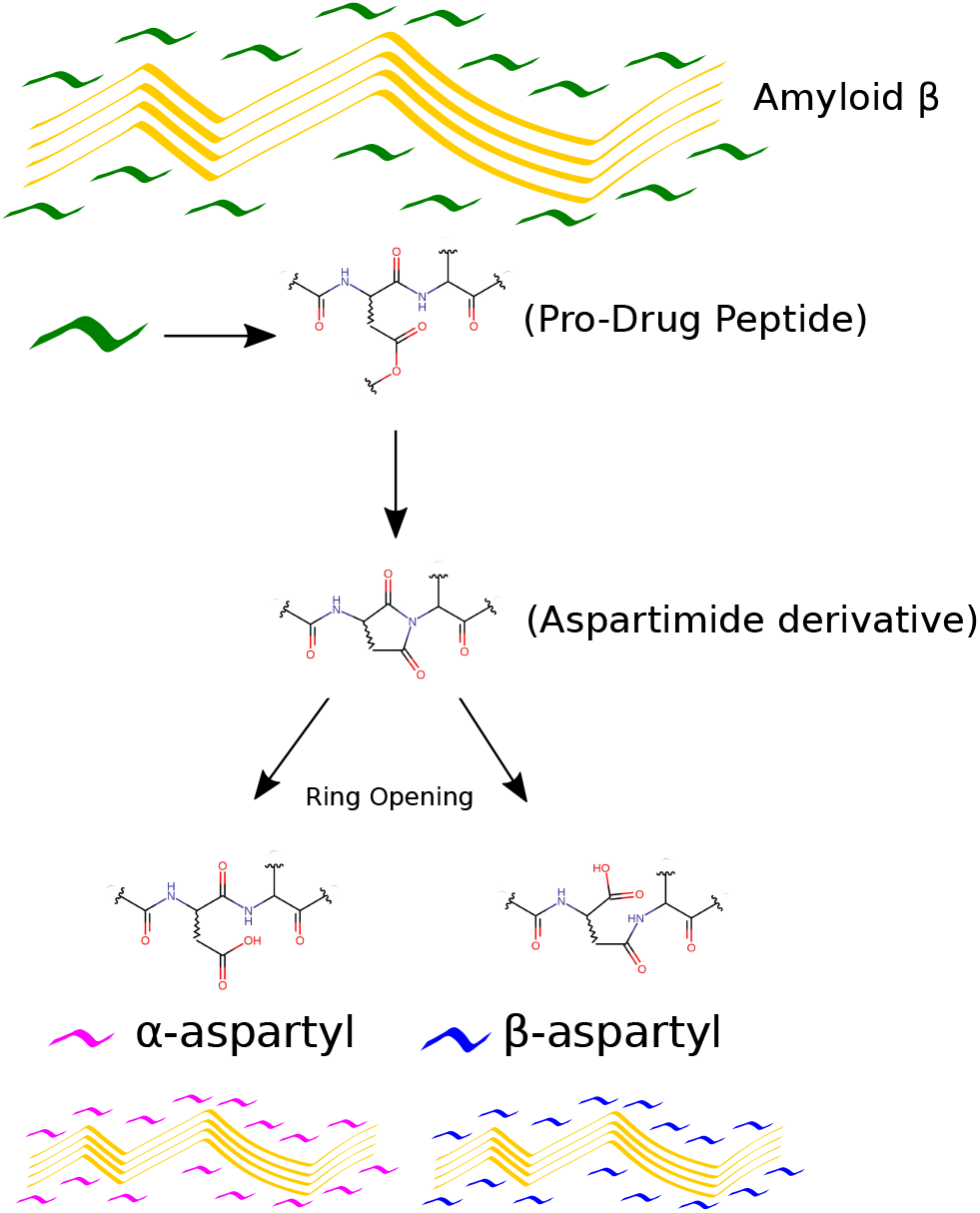
Conversion of Pro-Drug peptide into its metabolites and random placement of metabolites around the A*β* aggregate.

## 1 Results and discussion

Initially, control simulations were performed to study the inherent stability of the A*β* aggregate. Simulations were done for 500 ns without any ligands present around the protein. It is observed that Asp^1^, Lys^28^ which form an interchain saltbridge were stable (figure S5), and we also observed a partial disruption of chain F and chain I from the aggregate since these subunits are not bound by any interchain saltbridges (figure S3). Snapshots of the aggregate at various instants were provided in supporting information (figure S4).

Now we present the results of the simulation study with Pro-Drug peptide (PDp) as a ligand. Percentage contacts of PDp with each subunit (figure 1a,1b) of the aggregate shows that the ligand has an affinity towards a part of the subunit rather than the whole. This is due to the structural similarity of the PDp molecule with A*β*^19-20^ (-F-F-) of the subunit. This recognition motif leads to the grouping of PDp molecules near Lys^16^ to Ala^21^ residues of chain E (figure 1i, S6) which increased the nonbonded interactions (figure 1d) as well as the number of hydrogen bonds (N_*HB*_) between chain E and PDp and decrease in N_*HB*_ between the chain E & its adjacent chain (chain C) (figure 1c). A decrease in the beta-sheet content of the chain E is also observed (figure 1e) showing the disruptive tendency of the ligand.

**Figure 1:**
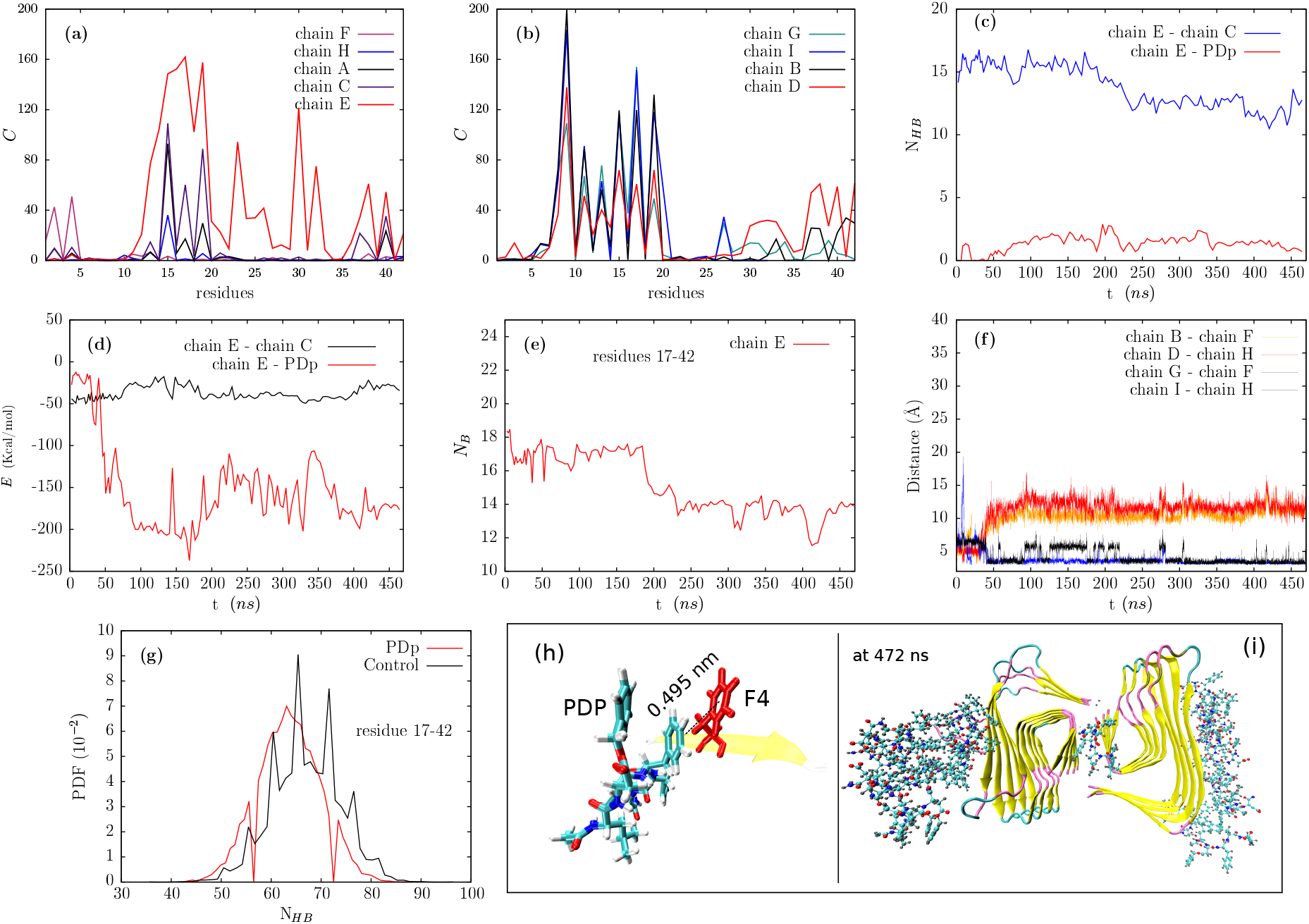
Simulation A2, Ligand: PDp, (a),(b) variation in percentage contacts (*C*) of PDp with residues of each subunit of A*β*, (c) number of hydrogen bonds (N_*HB*_) vs time (ns), (d) variation of nonbonded interactions (*E*) vs time (ns), (e) number of backbone carbons (*N_B_*) (which contribute to *β* sheet) vs time, (f) distance between the saltbridge composing atoms vs time, (g) probability density function (PDF) of number of hydrogen bonds (*N_HB_*) compared between PDp and control simulations, (h) snapshot showing the π-interactions between aromatic ring (A.r) of Phe^4^ of the subunit and the aromatic ring of the ligand with an observed angle of 10.4 degree between the aromatic planes and (i) snapshot of the clustering of PDp molecules near the chain E of A*β* aggregate.

We noticed that the terminal chain, i.e., chain F, partially came out from its natural position, which is considered usual as it is observed in control simulations, and the cause for the disruption is explained earlier. Moreover, we observed a slight disruption in one of the saltbridge in the presence of PDp. Initially, one of the aromatic rings of PDp have a preference in π-interaction, with the aromatic ring of the residue Phe^4^ (figure 1h) of chain D which pulled the chain out, but this interaction did not last for a long time and is due to the absence of a firm attachment that holds a PDp molecule to the terminal chain.

From figure 1f, we can infer that the increase in the distance of interchain saltbridge corresponds to the partial breakage of saltbridge between the chains B-F & chains D-H and decrease in the distance refers to the formation of new saltbridges between the chains I-G & chains G-F which holds the aggregate intact. Probability Density Function (PDF) of the number of hydrogen bonds compared for control and with PDp as a ligand shows that there is a reduction in number in the presence of PDp, i.e., peak value decreased from 66 to 61, indicating the disruptive tendency of PDp molecules (figure 1g).

Hydrolysis of aspartimide derivative produces racemized aspartyl derivatives (*α*/*β* 1:3)^*16*^ (figure S1, S2). Results for the *α*-aspartyl simulations are presented here. The *α*-aspartyl derivative also has a recognition motif with A*β*^19-20^ (-F-F), which helped these ligands to be drawn towards Phe^4^ (-F-) of chain D (the terminal chain) and is a crucial event for the breakage of interchain saltbridges. Initially, the α-aspartyl derivative was involved in simultaneous hydrogen bonding with Arg^5^ of chain D & chain B (figure 2g). The electron-rich aromatic rings and the acetyl oxygen of the ligand were involved in nonbonded interactions with electron cloud around NHCO of Phe^4^ of A*β*. These interactions are repulsive (figure 2e,2f), which forced chains B, D, I, and G to move away from these ligands (figure 2d), ultimately breaking their respective interchain saltbridges. We noticed that the hydrogen bonding between a ligand and Arg^5^ is functioning as a firm support, which allows the ligand to stay for a extended period in the vicinity of the terminal chain. From figure 2h and 2f, we can deduce that when the distance between the aromatic ring of the ligand and oxygen of Phe^4^ is less than 10 Å, there is a slight increase in the distance between the saltbridge forming residues which confirms that the saltbridge breakage is a consequence of this repulsion (figure 2d). The reason for the slight increase in the distance, i.e., slight breakage rather than complete breakage, despite the repulsions is due to the formation of new intrachain hydrogen bonds and will be explored in-depth in the upcoming paragraphs.

**Figure 2:**
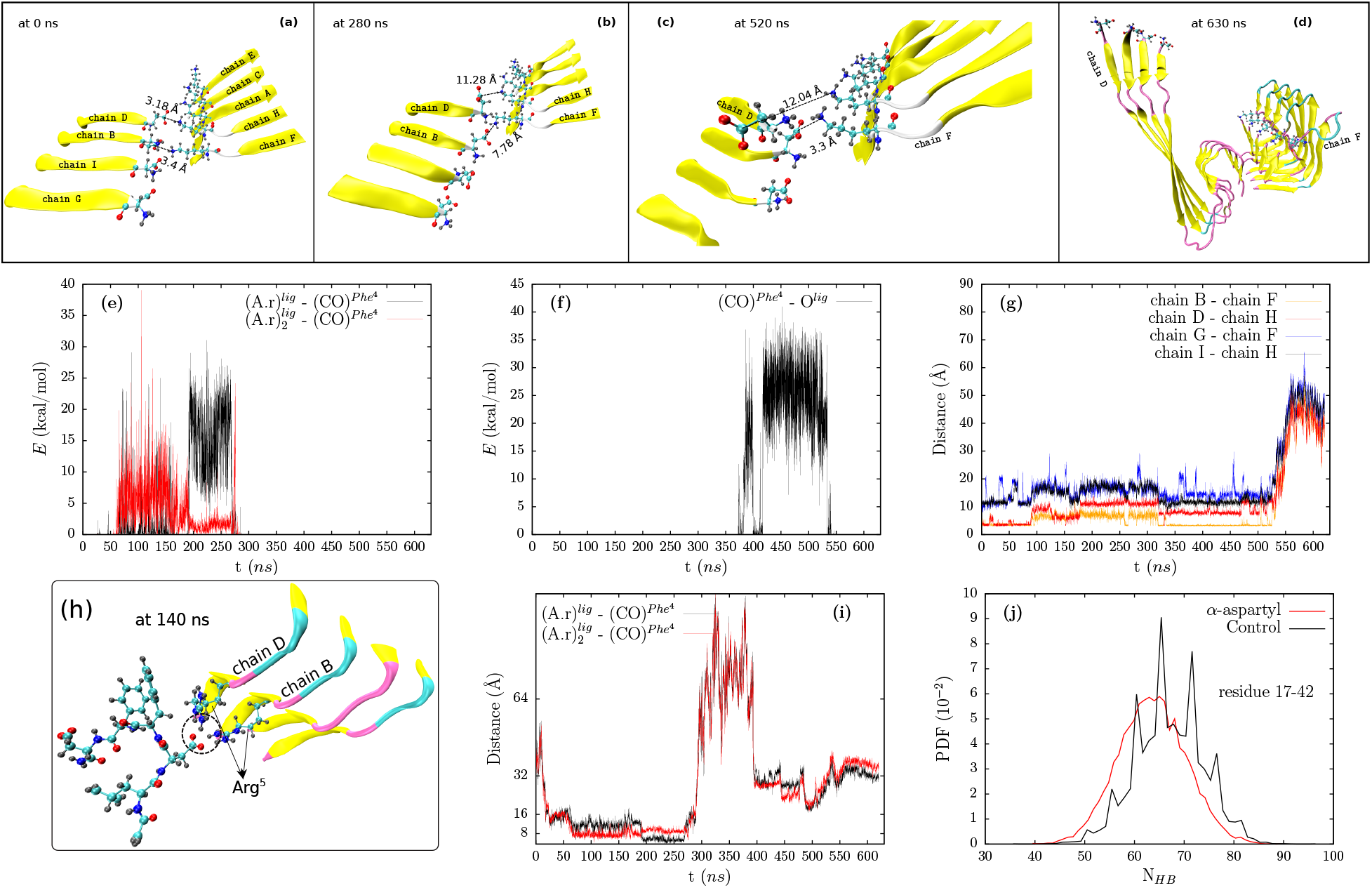
Simulation A3, Ligand: *α*-aspartyl, (a,b,c,d) snapshot showing the distance between the residues composing the saltbridge at 0, 280, 520 and 630 ns, variation in nonbonded interactions (E) with time between (e) aromatic rings of the ligand 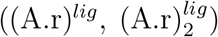 and the carboxyl oxygen of Phe^4^ ((CO)^*Phe*^4^^) of A*β*, (f) carboxyl oxygen of Phe^4^ ((CO)^*Phe*^4^^) of A*β* and the acetyl oxygen of the ligand (O^*lig*^), (g) variation in distance with time between the residues Asp^1^ and Lys^28^ which compose the salt bridges, (h) snapshot showing the hydrogen bonds formation (dotted circle) between the ligand and Arg^5^ of terminal chain, (i) variation of distance with time between oxygen of Phe^4^ of chain D of A*β* and aromatic rings of the ligand and (j) PDF of the number of hydrogen bonds present in the A*β* protofibril with and without *α*-aspartyl

From the figures 2a, 2b and 2c, we notice that the alignment of oxygen in the carboxyl group of Asp^1^ in chain D and nitrogen in the NH_2_ group of Lys^28^ in chain H is lost and did not regain for the rest of the simulation. In other words, carboxyl group and the NH_2_ group in D^1^ of chain D exchanged there positions, ultimately creating a repulsive interaction between the saltbridge forming residues. Due to this repulsion, the saltbridge between chain D-H did not recover fully whereas in the case of chain B-F recovery is seen after the ligand left the vicinity of chain D (decrease in distance and increase in attraction in figure 2g & figure S12 respectively between 320 & 520 ns). The main cause for this misalignment is due to the direct action of a-aspartyl on chain D, which leads Asp^1^ of chain D to form new hydrogen bonds with Gly^38^ of chain G (figure S9).

In figure 2g, we notice that the distance increased slightly after 170 ns and remained constant till 320 ns despite the repulsions between the aromatic ring of the ligand and the oxygen of Phe^4^ (figure 2e) which is believed to be the causative agent for the breakage of saltbridges earlier. This slight increase and maintaining constant distance is a consequence of the formation of the new intrachain hydrogen bond between Asp^1^ and Gly^38^, which stopped the chains further movement that can lead to complete breakage of saltbridges. However, after 530 ns we observe a drastic increase in the distance (figure 2g), this is due to the repulsion caused between the acetyl oxygen of the ligand and the carboxyl oxygen of Phe^4^ of A*β* (figure 2f, S11) which forced chains to move and ultimately breaking all the interchain saltbridges. This repulsion is so well aligned and strong that it also broke the newly formed hydrogen bond, which halted the movement of the chains earlier. The role of alignment of the ligands with the terminal chain in breaking the saltbridge will be discussed in the latter part of this article.

As a result of the interchain saltbridge breakage, the N-terminal part of one of the protofilaments which are considered to have a substantial influence on A*β* aggregation^*21*^ is entirely free from the other (Figure 2d, S7, S8, and S9). Concurrently, the hydrophobic cluster formed by the residues Leu^17^, Val^19^, and Ile^31^, which is crucial for the structural stability of the protofilament is also being gradually disrupted. As per Berhanu *et al.^22^* the interchain saltbridges are comparitively stronger than the intrachain saltbridges and they also demonstrated the importance of inter-sheet side chain-side chain contacts, hydrophobic contacts among the strands, saltbridges in the stabilization of A*β*.

The other metabolite formed after the hydrolysis of the aspartimide derivative is the *β*-aspartyl derivative. Simulations were performed using this as a ligand for 630 ns, and the results are presented here. The *α* & *β*-aspartyl derivatives are constitutional isomers. The backbone of the *β*-aspartyl is broken, which makes it behave as two flexible moving parts joined at one of the nitrogen atoms (figure S2 marked by*), whereas the *α*-aspartyl derivative and the PDp have a straight native backbone. We noticed that one of the two parts of the *β*-aspartyl bound firmly to one chain of A*β* while the other part attaches and detaches intermittently to another adjacent chain. This is well illustrated in figure 3h, where the binding part of the ligand has constant energy, and the flexible part has significant fluctuations in *E*. Due to this unique feature, the *β*-aspartyl ligand has more number of hydrogen bonds formed with subunits compared to the *α*-aspartyl (figure 3f).

**Figure 3:**
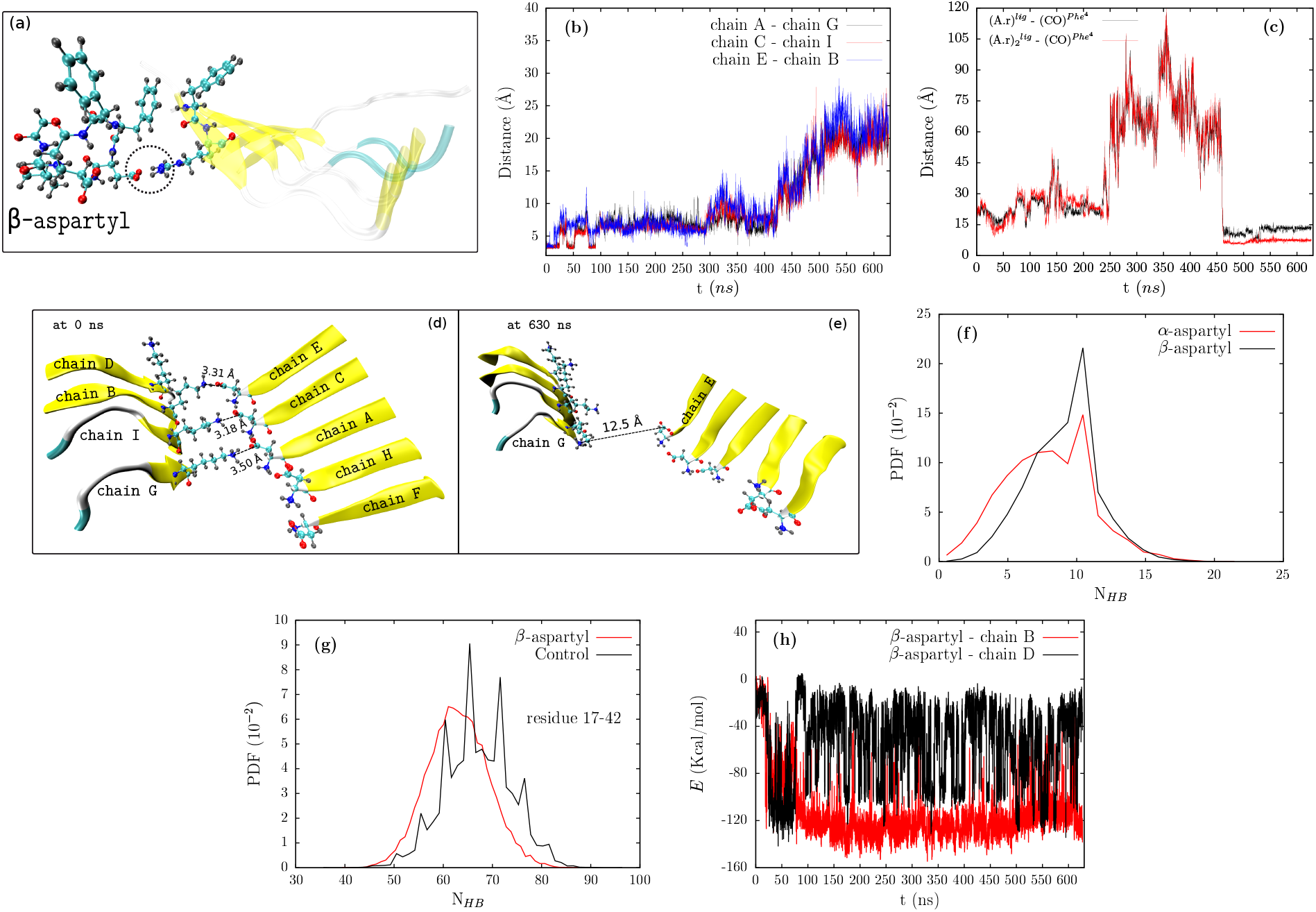
Simulation A4, Ligand: *β*-aspartyl, (a) snapshot showing the hydrogen bonds (dotted circle) between ligand and Arg^5^ of terminal chain, (b) variation of distance between the residues Asp^1^ and Lys^28^ which compose the saltbridges, (c) distance between the center of the aromatic rings of the ligand 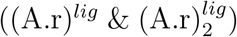 and carboxyl oxygen of Phe^4^ ((CO)^*Phe*^4^^), (d) & (e) snapshots showing the distance between the saltbridges, (f) comparision of PDF of number of hydrogen bonds formed between the ligand and the protein, (g) comparision of PDF of hydrogen bonds in the presence of ligand and control simulation and (h) variation of nonbonded interactions between the ligand and the chain B and chain D

Similar to the a-aspartyl ligand, we observed breakage of the interchain saltbridges in case of *β*-aspartyl (figure 3b, S20). The *β*-aspartyl was involved in hydrogen bonding with the residue Arg^5^ of the terminal chain (figure 3a), and the aromatic ring of the ligand was involved in repulsive interactions with backbone carboxyl oxygen of Phe^4^ of the same terminal chain (figure S19). Similar to the Simulation A3, breakage happened when the aromatic ring of the ligand is within 10 Å of the terminal chain (figure 3c).

In both simulations A3 & A4, initially, the saltbridge breakage happened through the repulsive interaction between the aromatic ring of the ligand and backbone carboxyl oxygen of Phe^4^. Although the distance between these repulsive groups is equal in both simulations (figure 2h, 3c), saltbridge breakage due to this repulsion is more pronounced in case of simulation A4 than A3. To understand this, we further investigated and found out that the orientation of the ligand when interacting with the terminal chain is the key to the disruption of saltbridges. In the previous case, *α*-aspartyl (simulation A3) attacked in such a way that the aromatic ring is directly below the terminal chain, which forced the chain to move up and lead terminal residue Asp^1^ to form hydrogen bonds with Gly^38^ of adjacent chains. Due to this, further breakage of saltbridge is stopped despite the repulsive interactions between the ligand and Phe^4^. Whereas, in the case of simulation A4, the aromatic ring attacked the terminal chain in a sideward fashion, which made the saltbridge breakage easy. Moreover, the backbone carboxyl oxygen of Phe^4^ of the terminal chain and the aromatic ring of the ligand should be perfectly aligned to observe the maximum breakage. Figures S17 shows favorable sideward alignment and unfavorable sideward alignment, which we observed in simulation A4. The above principles will also apply for the repulsions between backone carboxyl oxygen of Phe^4^ and the acetyl oxygen of the ligand (figure S11) by which we observed a complete breakage of saltbridges in case of *α*-aspartyl as a ligand, i.e., simulation A3 but not in A4.

The process of breakage of saltbridge is halted after 530 ns (figure 4b) by an unfavorable sideward alignment of the ligand, which lead to the formation of an extra hydrogen bond between Asp^1^ and Gly^38^ identical to the one we observed in simulation A3. Variation of nonbonded interactions, which are essential for the breakage of the above-discussed saltbridges, are provided in supporting information (figure S19). In all cases which involve saltbridge disruption, there is also a disruption of the intrachain hydrophobic cluster ^*23*^ formed by Ala^2^, Val^36^, Phe^4^ & Leu^34^ (figure S10, S17). Moreover, we observed buckling of the protofilament due to the intense pulling of residues His^13^ to Phe^20^ by the *β*-aspartyl towards the solvent (figure S15), which we did not observe in any of the control simulations. Based on the existing simulation data, we propose that the action of ligands on His^13^ to Phe^20^, result in disruption of hydrophobic cluster^*23*^ formed by Leu^17^, Ile^31^ & Phe^19^, which is critical for the structural stability of the protofilament. When such disruption of the hydrophobic cluster is combined with the disruption of saltbridge, as we observed in simulation A3, we can expect a complete breakage of the protofilament. Figure 3g shows the PDF of the number of hydrogen bonds compared with the control simulation.

**Figure 4:**
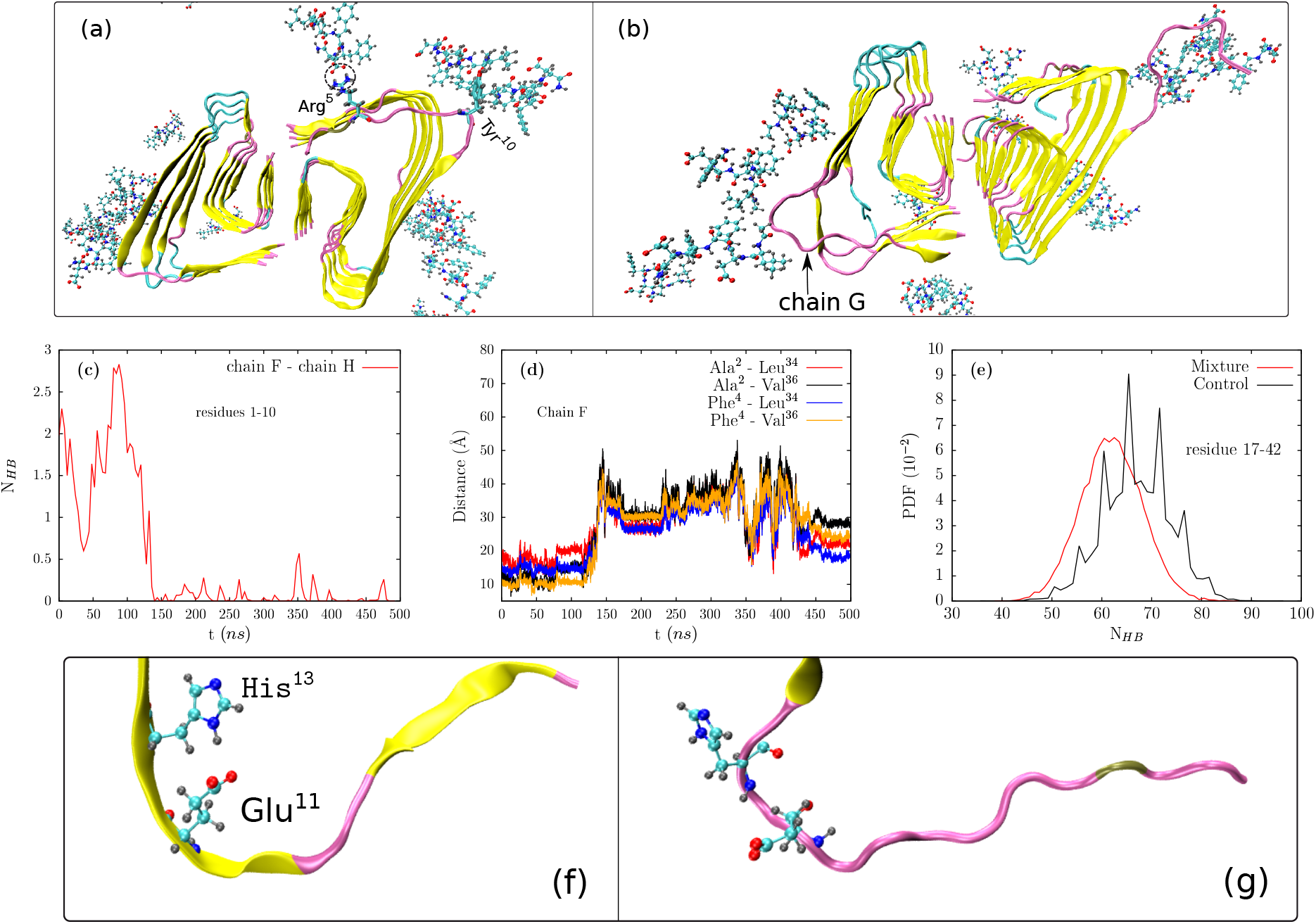
Simulation: A5, Ligand: mixture of *α* & *β* aspartyl in 1:3 ratio, (a,b) snapshot showing the modes of action through which the terminal chains disrupt in ligand environment, residues Arg^5^ & Tyr^10^ of chain F is shown in licorice representation, (c) variation in number of hydrogen bonds (N_*HB*_) vs time (t), (d) variation in distance between the residues forming the intrachain saltbridge vs time, (e) PDF of number of hydrogen bonds of the protein compared between control simulation and simulation A5, and (f, g) snapshot showing the saltbridge between Glu^11^ and His^13^ of chain G and its disruption in later part of the simulation due to the action of ligands.

A mixture of the *α* & *β* aspartyl derivatives in 1:3 ratio (similar to the experimental ratio) were used as ligands and simulated for 500 ns. As stated earlier, partial disruption of terminal chains is possible and is observed in control simulations. Nevertheless, on the addition of ligands to the system, the terminal chains experience less solvent accessibility, and in some instances, these ligands can obstruct the terminal chain to come out. Despite this obstruction and less solvent accessibility, partial disruption of the terminal chain is observed and is solely due to the interaction of the ligands with these chains. We noticed two modes of interaction by the ligand, which are responsible for the disruption of terminal chains. i)

The first mode of interaction is by formation of hydrogen bonds with Arg^5^ (figure 4a marked by a dotted circle) and the nonbonded interaction with Tyr^10^ by the ligand simultaneously. These concurrent interactions result in pulling the chain out towards the solvent which leads to the disruption of interchain hydrogen bonding between the chain that has been pulled out and its adjacent one (figure 4c) as well as the disruption of the hydrophobic cluster formed by Ala^2^, Val^36^, Phe^4^, Leu^34^ (figure 4d). ii)The second mode is by the ligand interactions with Val^12^, His^14^ (chain G of figure 4b) resulted in breakage of saltbridge between Glu^11^ and His^13^ (figure 4f, 4g) which is the key for the stabilization of kink in the N-terminal part of the *β* sheet around Y10^*23*^ of A*β* strand. Moreover, we did not observe any breakage of interchain saltbridges composed by the residues D^1^ & K^28^. Figure 4e shows that there is a decrease in the number of hydrogen bonds compared to control simulations. Although in the simulation with mixture of ligands, we did not observe the same that observed in the case with individual ligands, this is probably due to the number of individual ligands were less in this case. but we can well assume that the phenomena observed in simulation A3, A4 can occur here too. Snapshots of the simulation A5 and variation in percentage contacts of mixture of ligands with residues of A*β* were provided in supporting information (figure S21.S22).

## 2 Secondary structure analysis and RMSD

Secondary structure calculations were done and percentage beta-sheet, turn and coil were reported in the figure 5. In case of beta-sheet content, A*β* aggregate at 0 ns have a beta-sheet content of 78.57%. Control simulations resulted in the decrease in the beta-sheet content to 63.4%. As in the simulations which include ligands PDp,alpha-aspartyl,beta-aspartyl the beta-sheet content observed is 64.82%, 65.93% and 65.81% respectively. The lower betasheet content in control simulations can be attributed to partial disruption of the terminal chains of Abeta aggregate which was not observed in other three. Meanwhile in case of simulation A5 (mixture of alpha and beta aspartyl used as ligands), a beta-sheet content of 59.84% is observed. Significant decrease in beta-sheet content can be explained by the partial disruption of the terminal chains and other effect of ligands on A*β*. Increase in the amount of coil also adds to the better inhibiting capability of the molecules.

**Figure 5:**
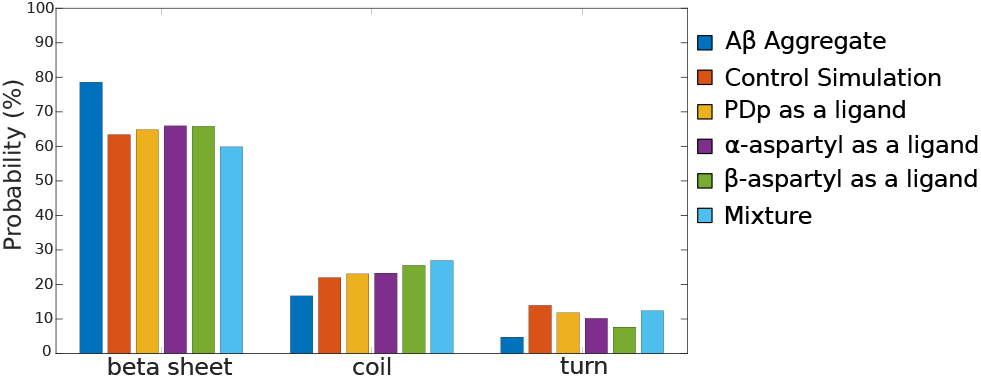
Average probability of each secondary structure contents of A*β* aggregate

Root Mean Square Deviation (RMSD) calculations for the protein backbone are shown in the figure 6. For the simulations where partial disruption of terminal chains observed, RMSD is calculated including and excluding those terminal chains. Due to the flexibility in the terminal chains by partial disruption, the RMSD values spiked over 5Å. But in the case of simulations which involve *α* & *β*-aspartyl as ligands, the spike in the RMSD is due to the breakage of the interchain saltbridges.

**Figure 6:**
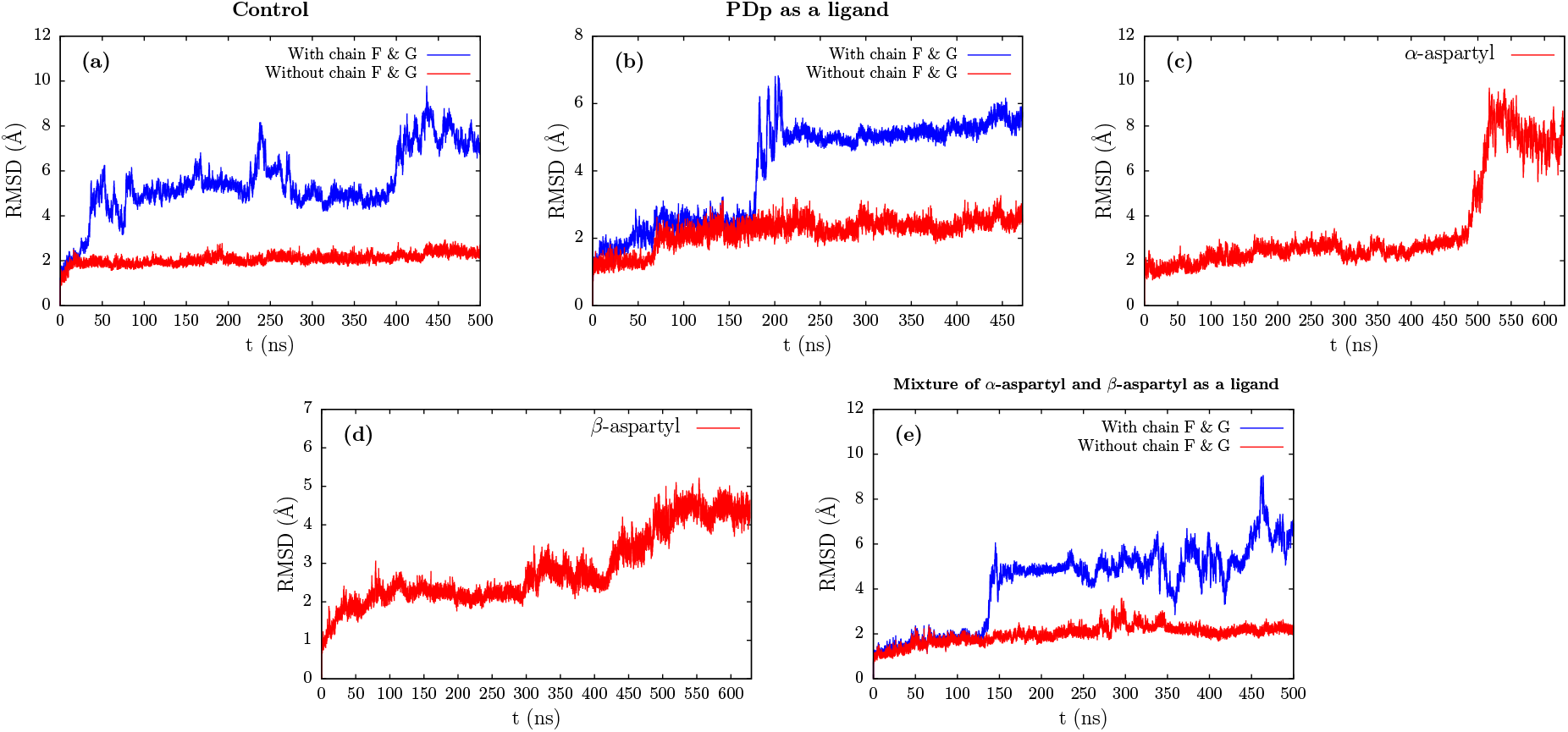
Variation of RMSD for the protein backbone with time

## 3 Conclusions

In summary, our work on the destabilization of A*β* aggregate using synthetic peptides provided a key insight into the mechanism of the fibril disruption by the Pro Drug peptide (PDp). PDp, which carries a sequence homology with a part of the A*β*, functions precisely as we intended by recognizing the A*β*^19-20^ (-F-F-) part of the protein and showing high affinity towards it. Notably, these PDp molecules are also capable of disrupting the amyloid aggregate. In case of the metabolites of PDp, *α* and *β* aspartyl as a ligand, one of the key findings is the formation of hydrogen bonds, which helps the ligand to attach to the terminal chain firmly and the repulsive interactions between Phe^4^ and ligand are mainly responsible for the disruption of saltbridges which in turn leads the N-terminal part of the protofilament to be entirely exposed to the solvent (Scheme 2). Moreover, by examining the *α* & *β*-aspartyl simulations, we understood that the orientation of the ligand with the terminal chain is crucial for the disruption of the saltbridges formed between Asp^1^ and Lys^28^. To the best of our knowledge, this is the first effort on simulation of a dynamic pro drug type ligand that not only acts as a *β*-sheet breaker peptide itself but also generates two metabolites that are more effective beta-sheet breakers. The present simulation study provided some crucial insights into the mechanism of destabilization of A*β* aggregate by PDp and its metabolites, hence we believe that this insights can help in designing a better drug for the disease. Eventhough in the present simulation study we considered PDp and its derivatives individually the dynamic evolution of the chemical reaction of PDp into its metabolites in computational framework will be explored in the future.

**Scheme 2:**
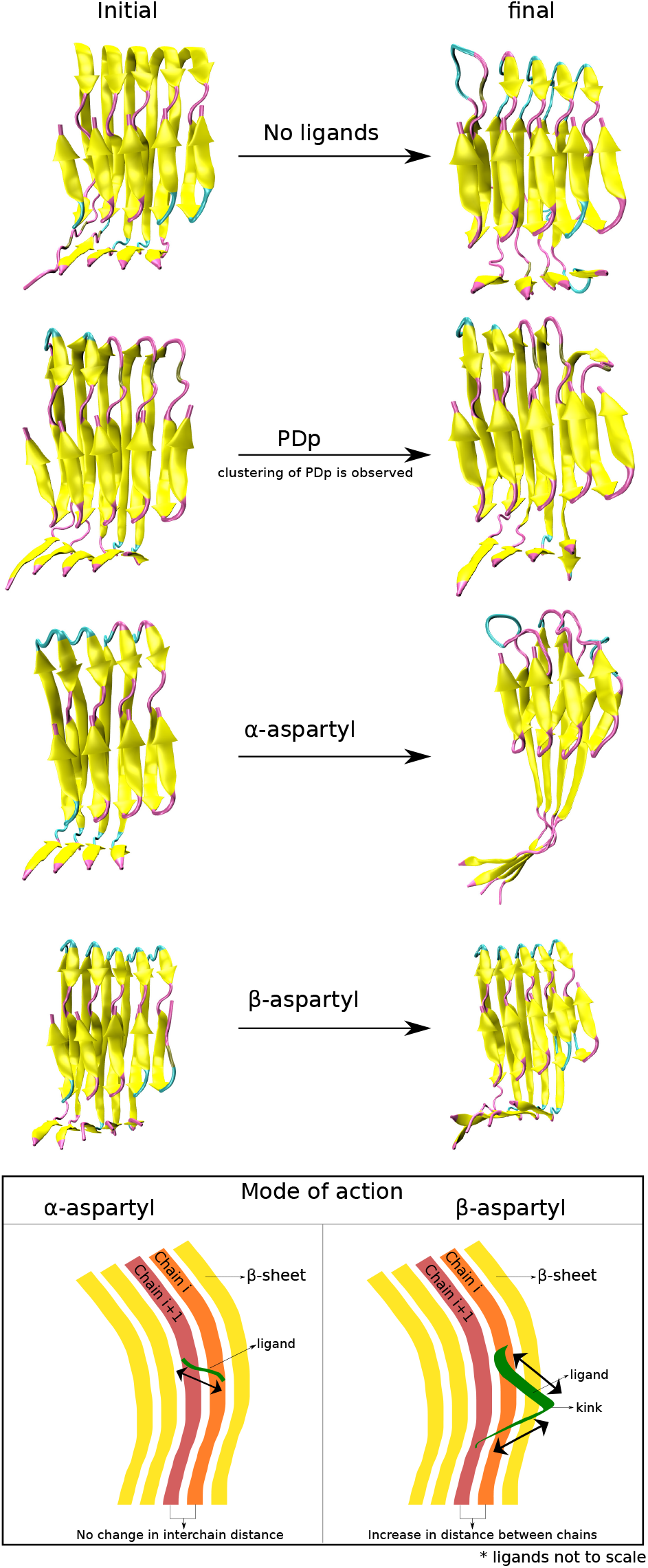
Initial and final snapshots of the A*β* protofilament in various environments and mode of action of metabolites of PDp with chains of A*β*. Double headed arrow represents the movement possible by the ligand.

## 4 Methodology

PDB ID 5OQV, an intertwined A*β* protofibril developed by Gremer *et al.^23^* is used as an initial structure for the simulations in the present study. This protein consists of 9 subunits, namely A B C D E F G H & I. Terminal residues (residues 1 to 5) of the subunits I G F & H are flexible since they are not bound by any saltbridges (figure S3). On the other hand, Pro Drug Peptides (PDp’s) and its metabolites were used as a ligand and were spread randomly around the protein. Eighteen ligands were used in each simulation setup maintaining a 2:1 ratio of ligand to the number of subunits of the A*β*. PDp’s are initially free from structure disrupting element but produces it in situ in the physiological conditions. PDp’s were converted to aspartimide derivative, which further hydrolyse into *α*-aspartyl & *β*-aspartyl. In the present study aspartimide derivative, due to the unavailability of the force field parameter and since it is a transient complex, we did not study its effect on A*β* aggregates, however, this will be taken up in the near future.

Classical molecular dynamics simulations were employed to investigate the destabilization of A*β* on the addition of the above-discussed peptides. For this purpose, five different systems were created using VMD^*24*^ version 1.9.3, namely A1, A2, A3, A4, A5. A1 is the control simulations where there are no ligands present, whereas systems A2, A3, A4 & A5 contains ligands, such as PDp, *α*-aspartyl, *β*-aspartyl & mixture of *α* & *β* aspartyl respectively. These simulations were performed using NAMD^*25*^ version 2.13 with CHARMM36^*26*^ as the force field for proteins, and the parameters for the ligand were obtained using CGENFF^*27,28*^ version 1.0.0. In each system, 18 ligands were added, and solvation is done using the TIP3P^*29*^ water model. 0.05 mol/l concentration of NaCl is added to replicate the physiological environment. Sufficient Na^+^ ions are added to neutralize the whole system. Long-range electrostatic interactions were computed using the particle mesh Ewald method.^*30*^ Van der Waals interactions are truncated at a cutoff distance of 1.2 nm, and the time step employed is 1 fs. In all cases, minimization is done for 5000 steps, and the equilibration run is for 2 ns in the NPT ensemble at 310 K, followed by production run in the NVT ensemble.

## Acknowledgement

The use of the HPC parallel computational facility PARAM-ISHAN at IIT Guwahati is greatly acknowledged.

## Supporting Information Available

Snapshots of each of the simulations at regular intervals of time and the ligands used in the present study, chemical reaction, information about the saltbridges, percentage contacts of ligands (*α*-aspartyl, *β*-aspartyl and mixture) with each of the subunit, buckling of the protofilament in case of simulation A4, favourable and unfavourable allignment of the ligand with the terminal chain, variation in distance of the hydrophobic cluster groups and the interaction energy between the groups which are responsible for the breakage of saltbridge.

## Notes

### Competing Interest Statement

The authors have declared no competing interest.

