## supporting information for "Pro-drug peptide and its metabolites disrupt amyloid fibrils by destabilizing salt bridge interaction and planar beta-sheet topology"

#### Supplementary Information

Table 1: Summary of the simulation time and number of atoms in a individual run

| System | Time (ns) | No. of Atoms |
|---|---|---|
| Control (A1) | 500 | 98419 |
| Pro Drug peptide (A2) | 472 | 133731 |
| $\alpha$-aspartyl (A3) | 630 | 146728 |
| $\beta$-aspartyl (A4) | 630 | 136289 |
| Mixture (A5) | 500 | 127150 |

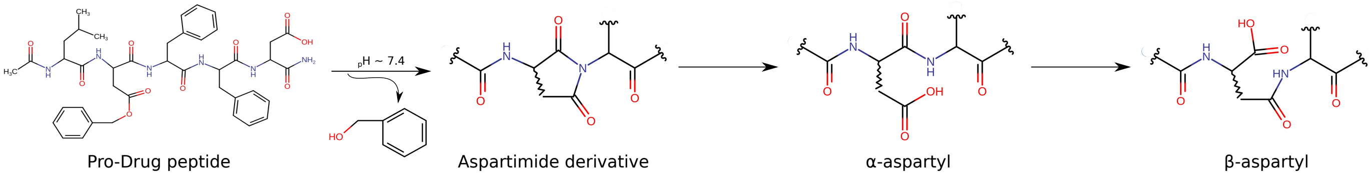


Figure S1: Chemical conversion of PDp and insitu generation of structure destablilizing elements at physiological conditions ( pH 7.4 and temperature 37°C )

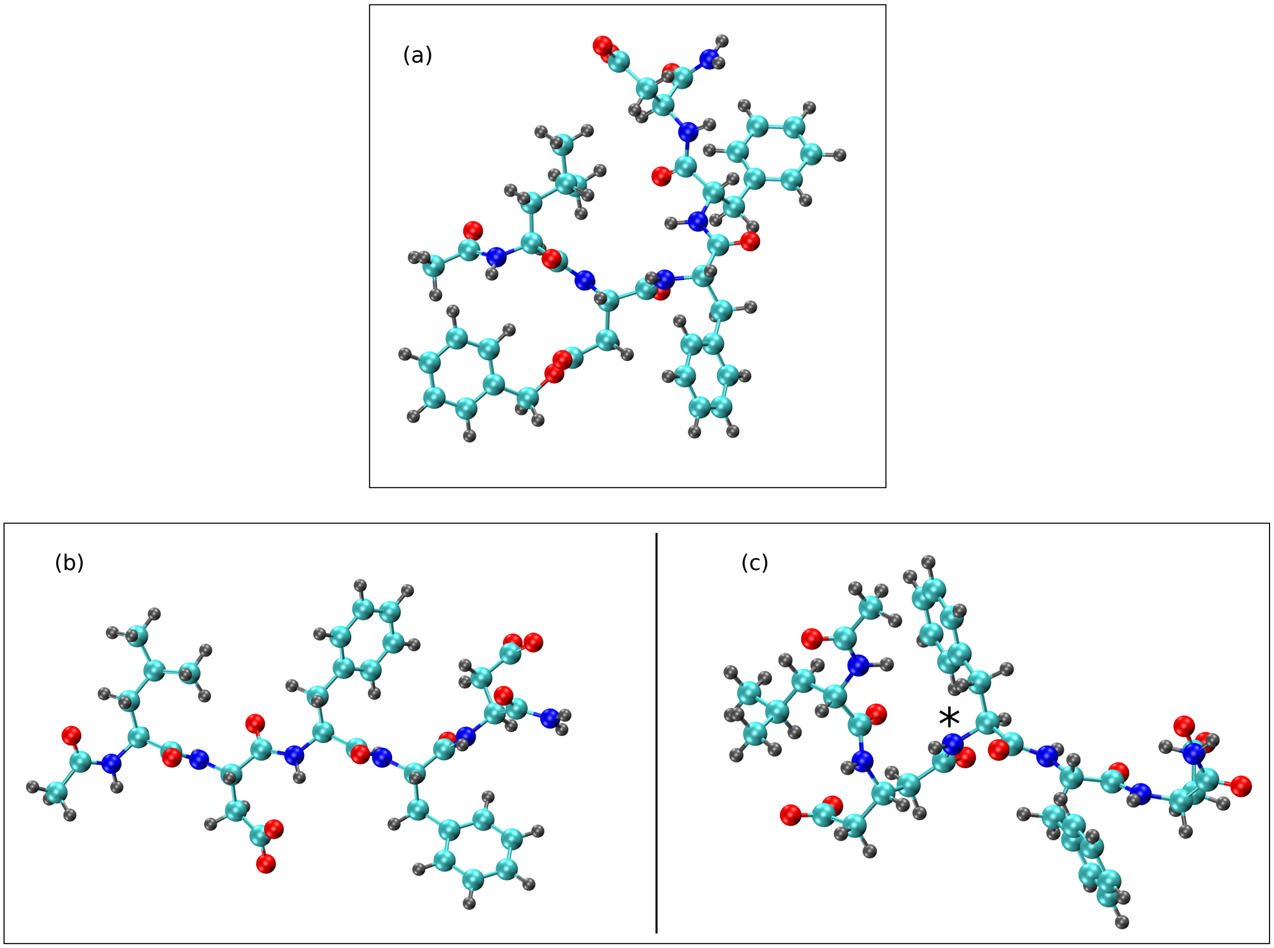


Figure S2: CPK representation of (a) Pro-Drug peptide (PDp), (b) $\alpha$-aspartyl and (c) $\beta$-aspartyl

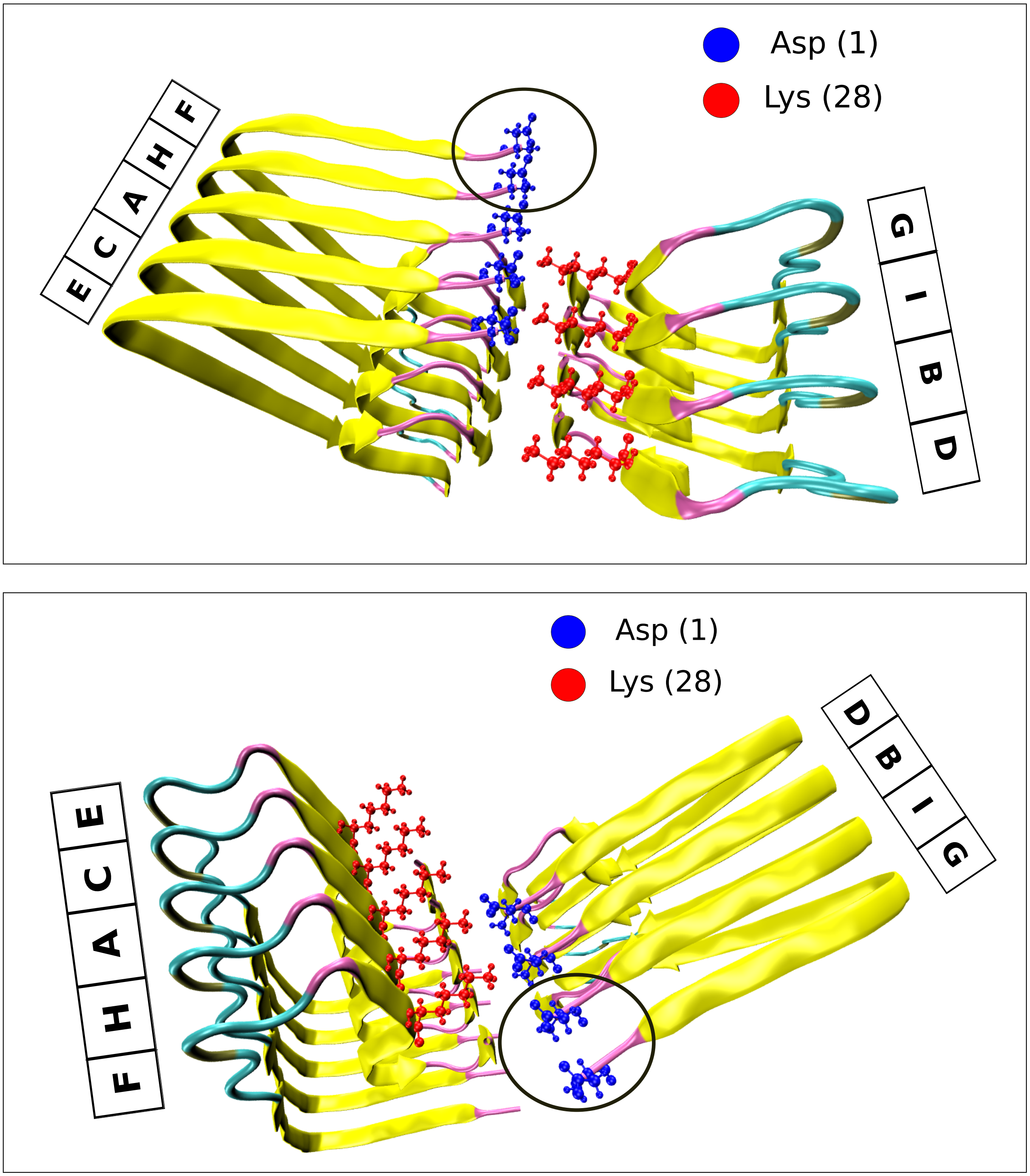


Figure S3: Snapshots showing the N-terminal part of the protofilament lacking interchain saltbridges shown by a black circle

### Simulation A1, Ligand: Control Simulation

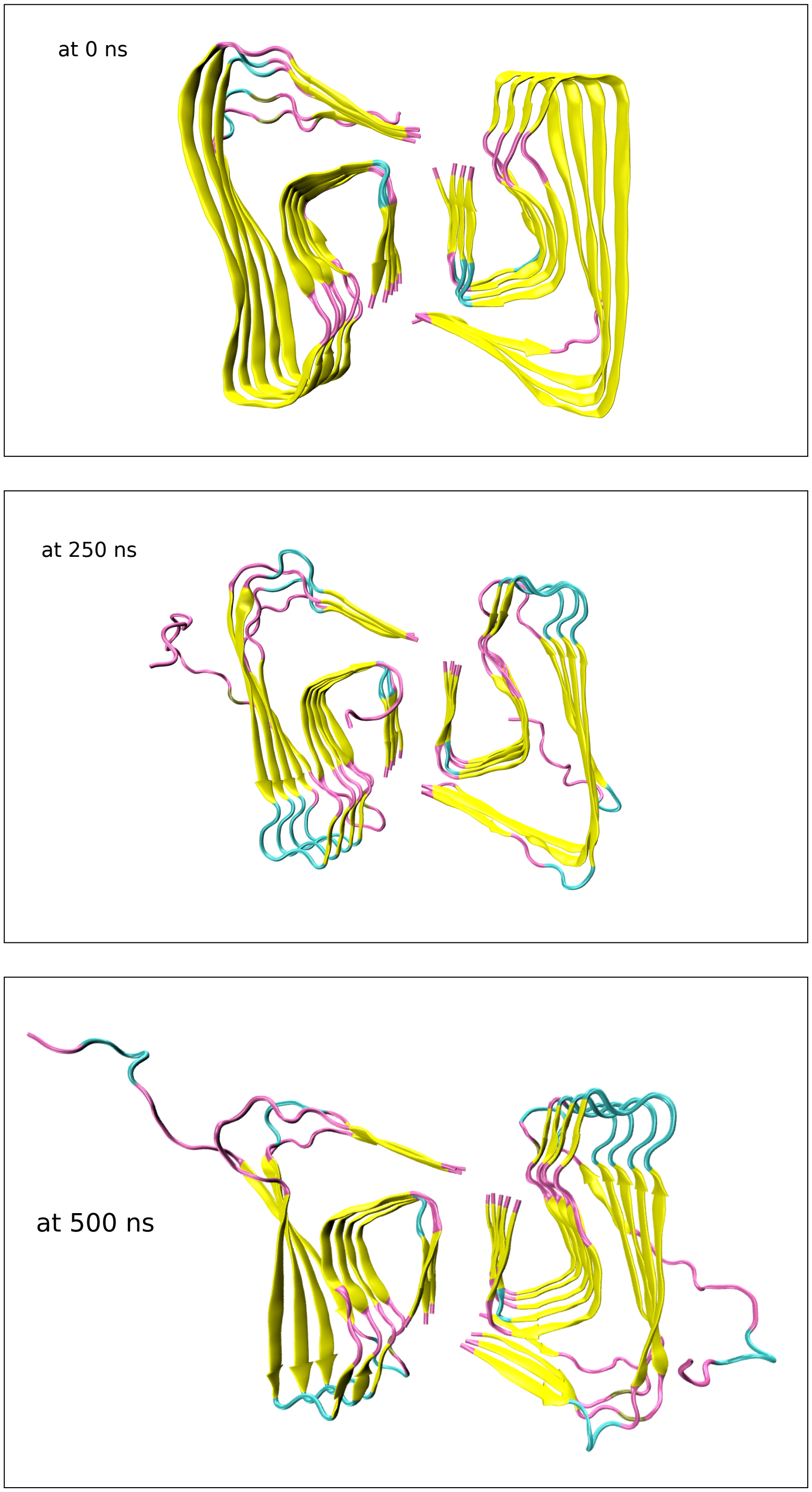


Figure S4: Snapshots of control simulation at 0, 250 and 500 ns. From 250 ns, we can see the N-terminal part of the chains F and G are free and exposed to the solvent due to the absence of interchain saltbriges.

### Simulation A1, Ligand: Control Simulation

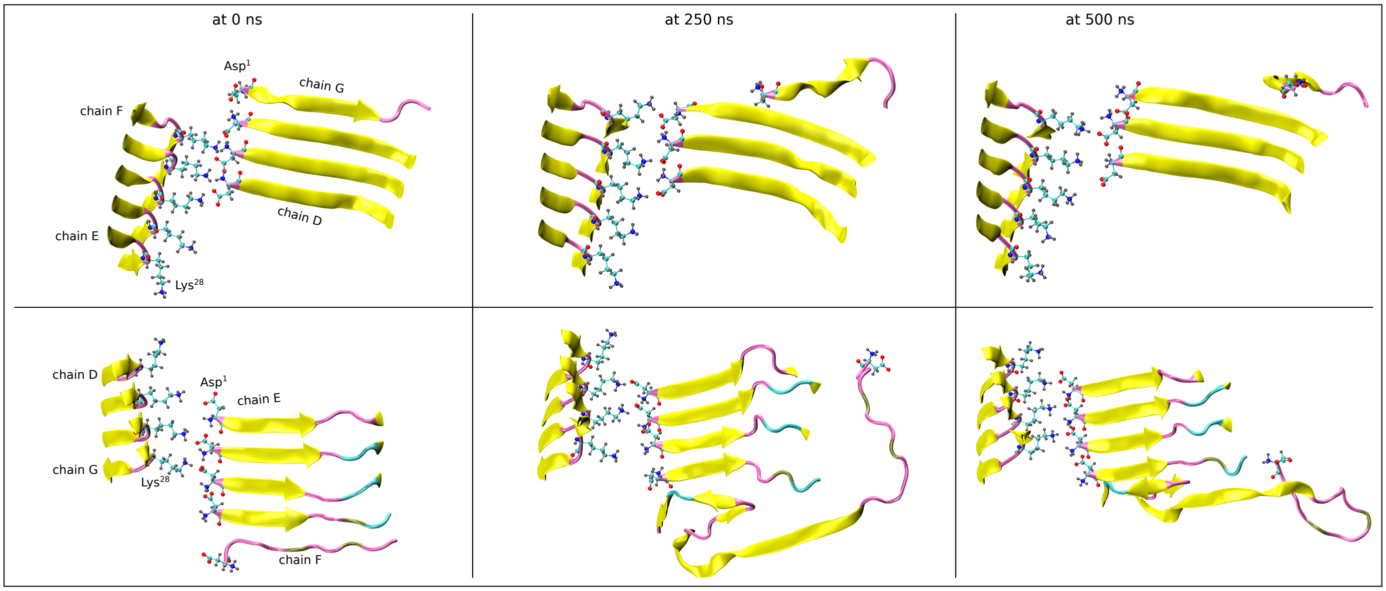


Figure S5: Snapshots of control simulation showing the interchain saltbridges between protofilaments at 0, 250 and 500 ns

### Simulation A2, Ligand: Pro-Drug peptide (PDp)

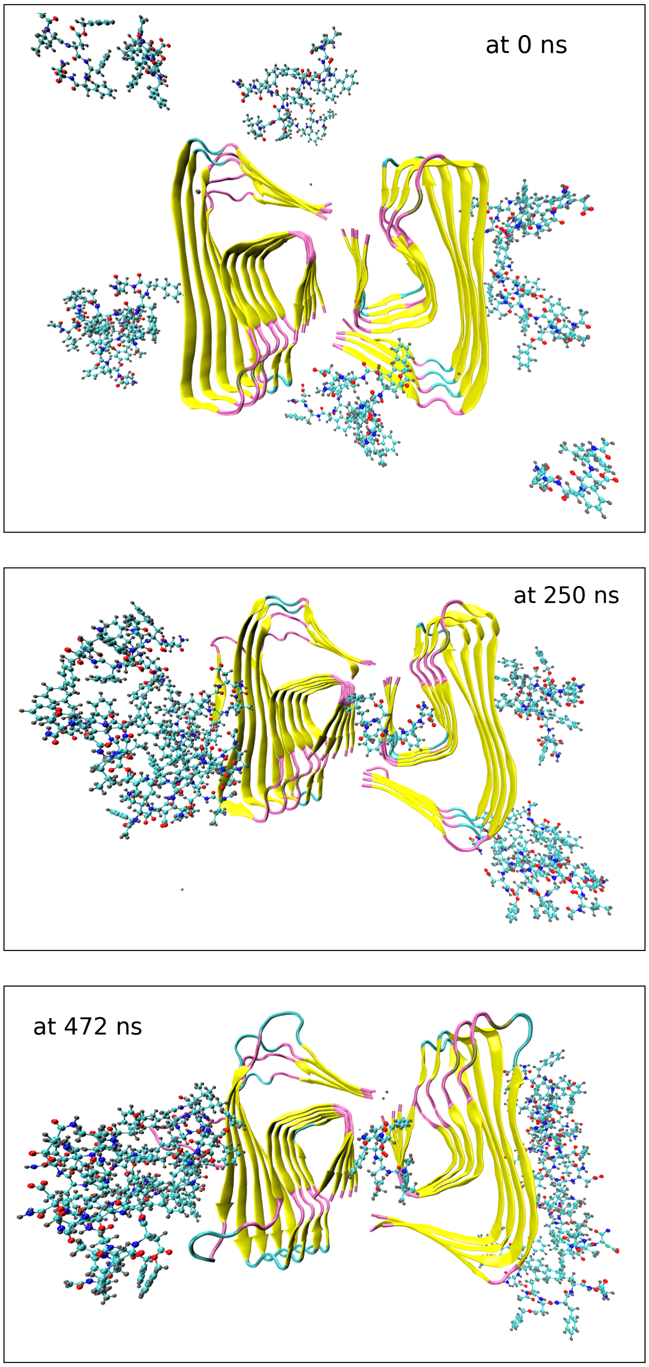


Figure S6: Snapshots showing the protein and surrounding PDp molecules at various instants of the simulation. The recognition motif carried by the PDp molecules is responsible for the clustering.

### Simulation A3, Ligand: $\alpha$-aspartyl

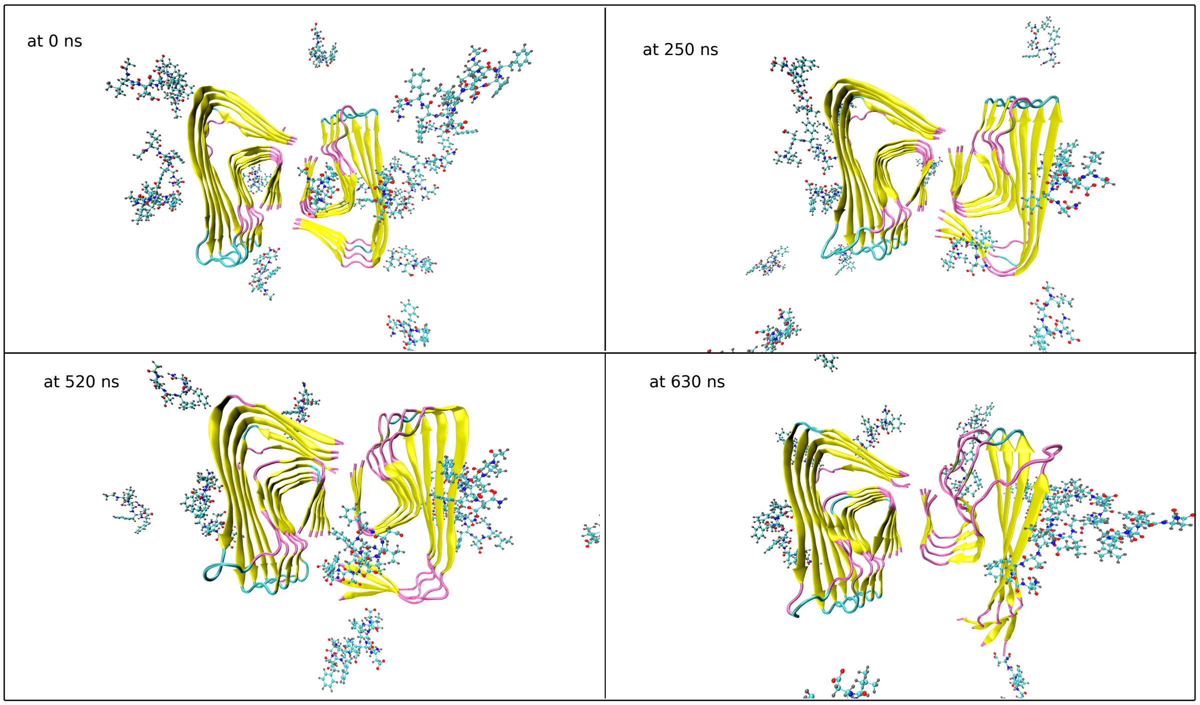


Figure S7: Snapshots from the simulation A3 at 0, 250, 520 and 630 ns

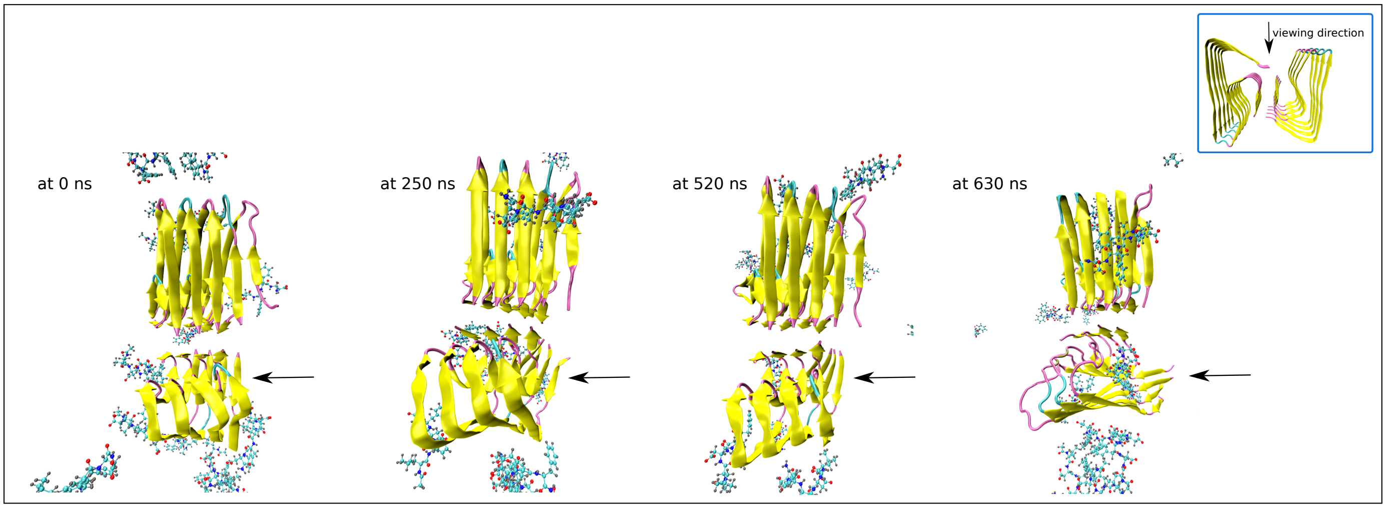


Figure S8: Snapshots showing the topview and the arrow pointing the effect of breakage of saltbridges

### Simulation A3, Ligand : $\alpha$-aspartyl

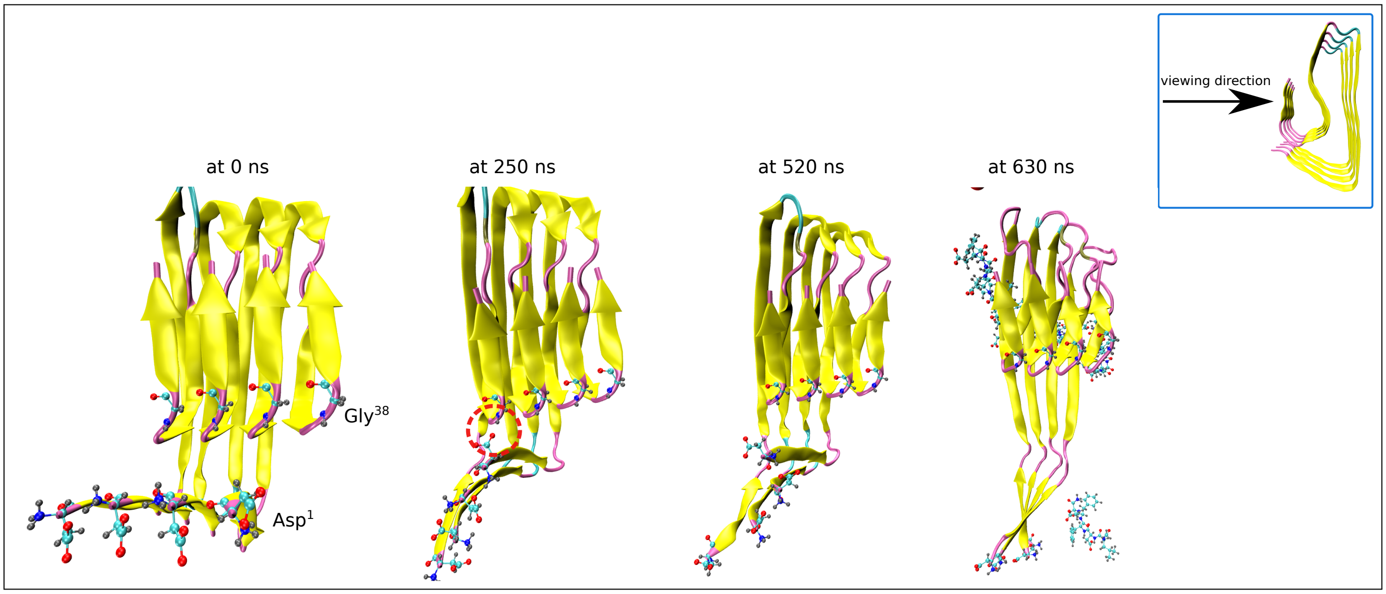


Figure S9: Snapshots showing the formation of new hydrogen bonds between Asp$^1$ and Gly$^{38}$ represented by red dotted circle

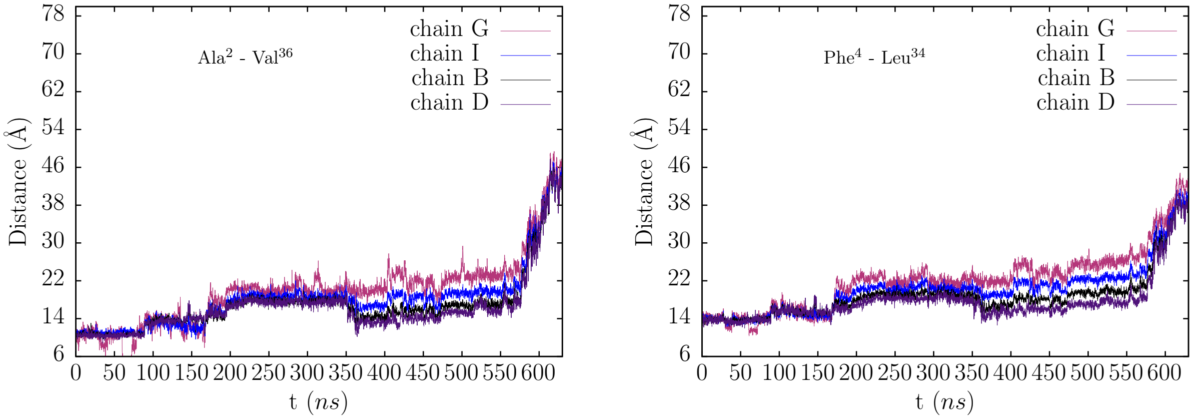


Figure S10: Variation in distance with time between the residues forming hydrophobic cluster.

### Simulation A3, Ligand: $\alpha$-aspartyl

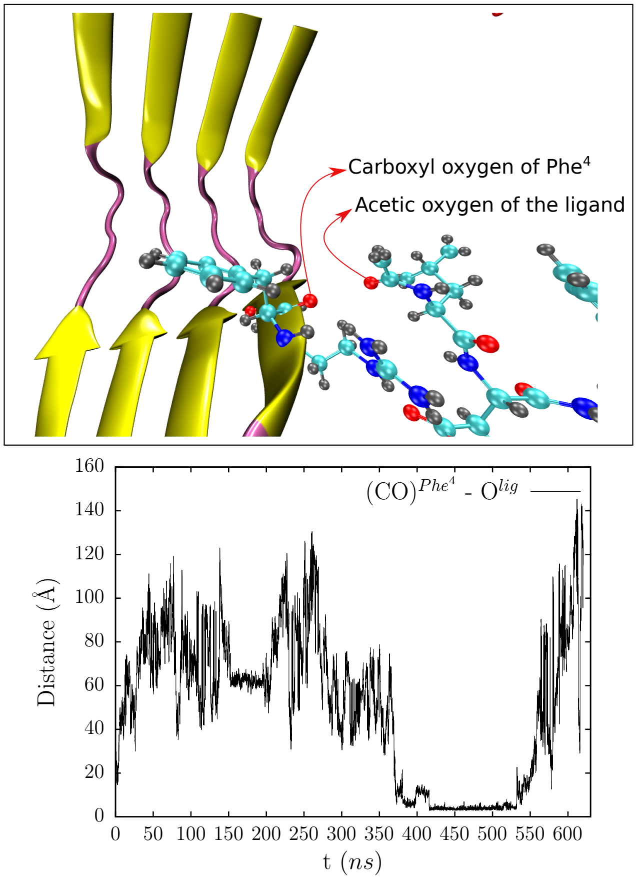


Figure S11: (a) Snapshot showing the carboxyl oxygen of residue $Phe^4$ of chain D and acetyl oxygen of the ligand, (b) variation of distance between carboxyl oxygen of $Phe^4$ ($(CO)^{Phe^4}$) and acetyl oxygen of the ligand ( $O^{lig}$ )

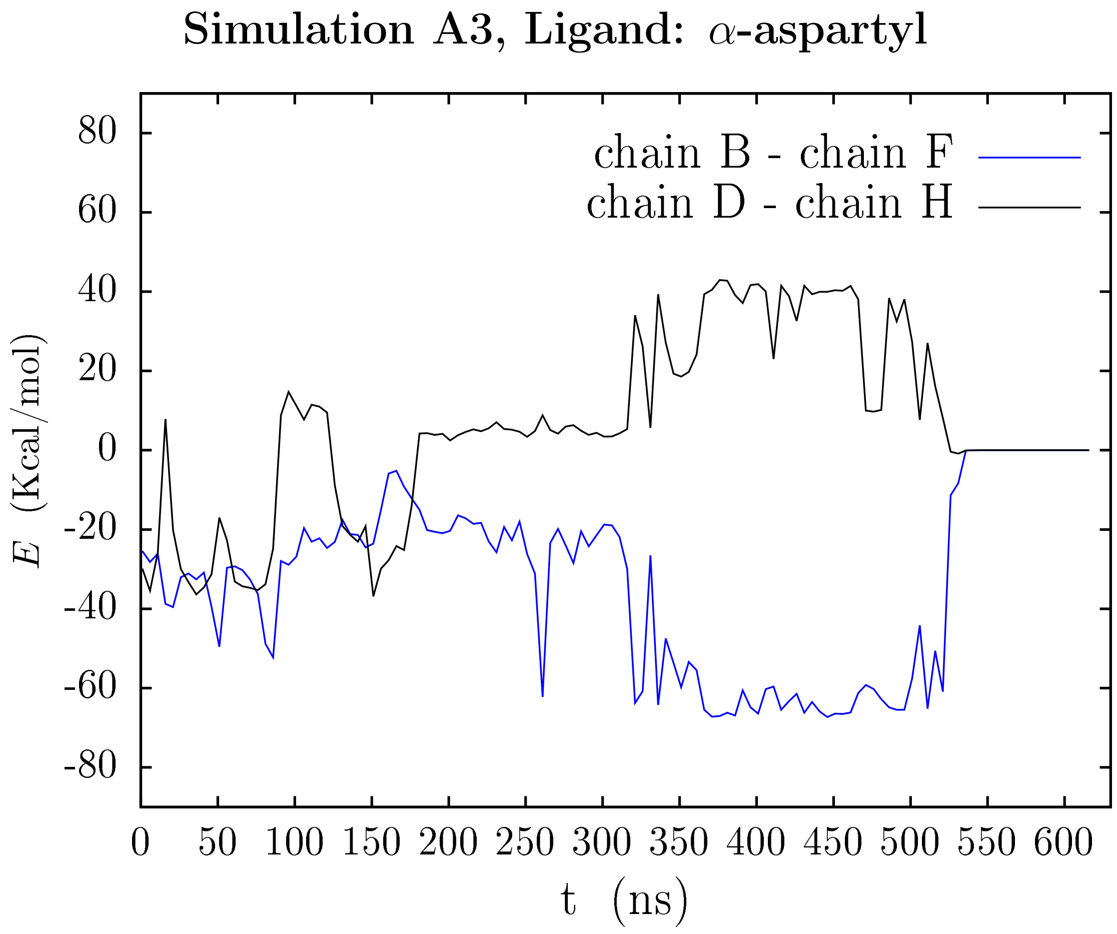


Figure S12: Variation of nonbonded interactions of interchain saltbridges composed by Asp[1] and Lys[28] of chain B-F and chain D-H.

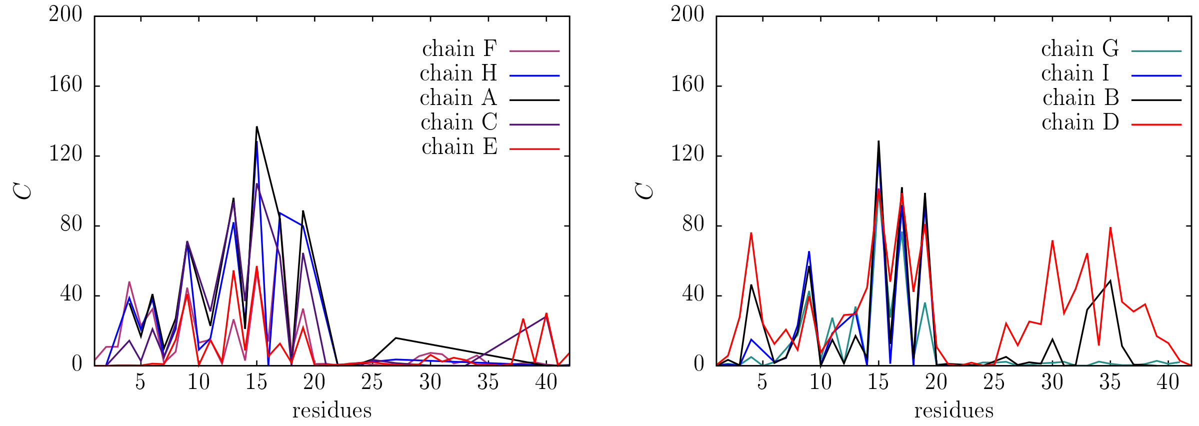


Figure S13: Variation in percentage contacts ($C$) of $\alpha$-aspartyl with residues of each subunit of A$\beta$

### Simulation A4, Ligand: $\beta$-aspartyl

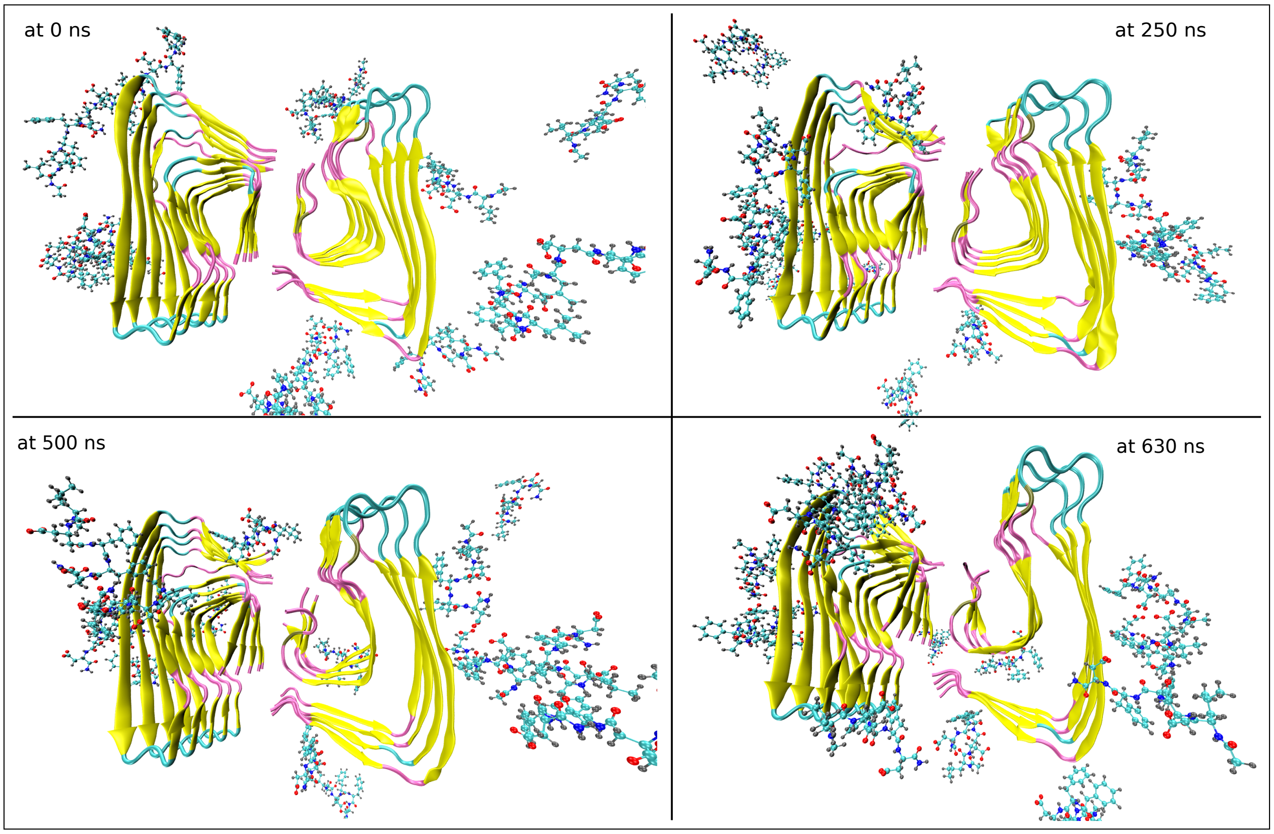


Figure S14: Snapshots for simulation A4

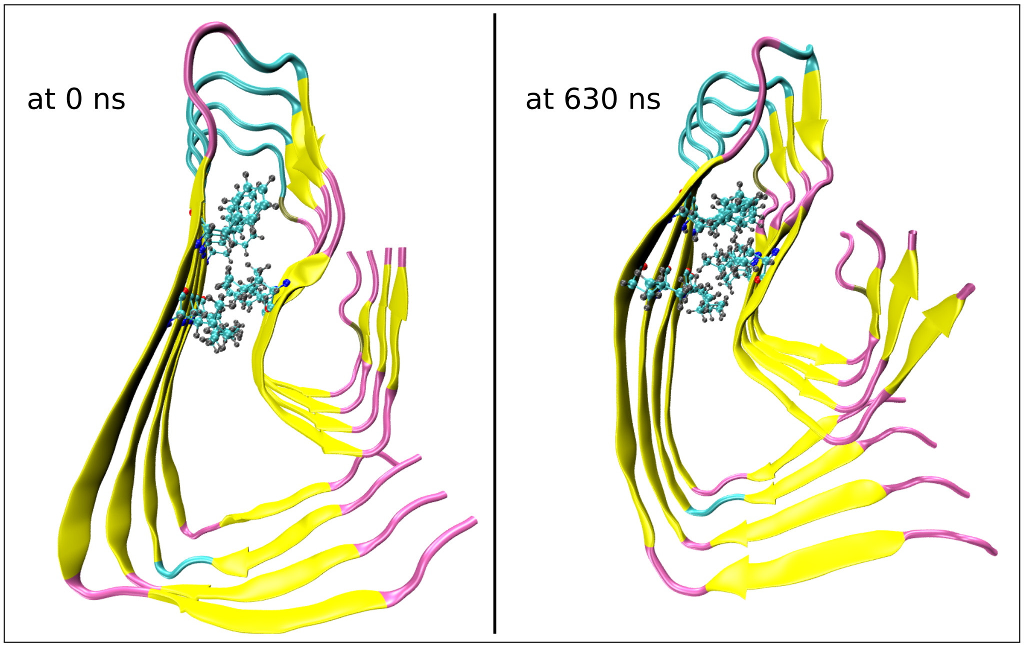


Figure S15: Snapshots showing the buckling of the fibril which might disrupt the hydrophobic cluster between $Leu^{17}$, $Ile^{31}$ and $Phe^{19}$ due to the action of ligands

### Simulation A4, Ligand: $\beta$-aspartyl

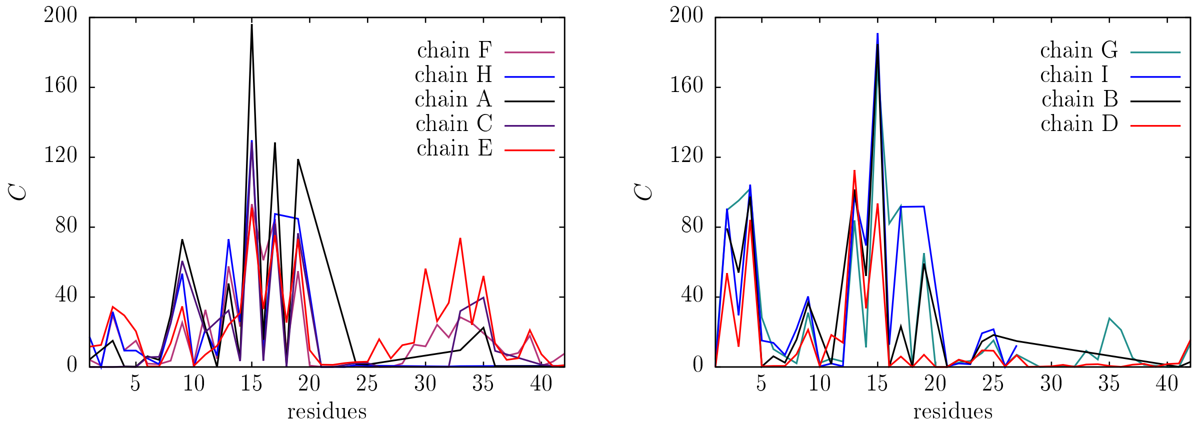


Figure S16: Variation in percentage contacts ($C$) of $\beta$-aspartyl with residues of each subunit of A$\beta$

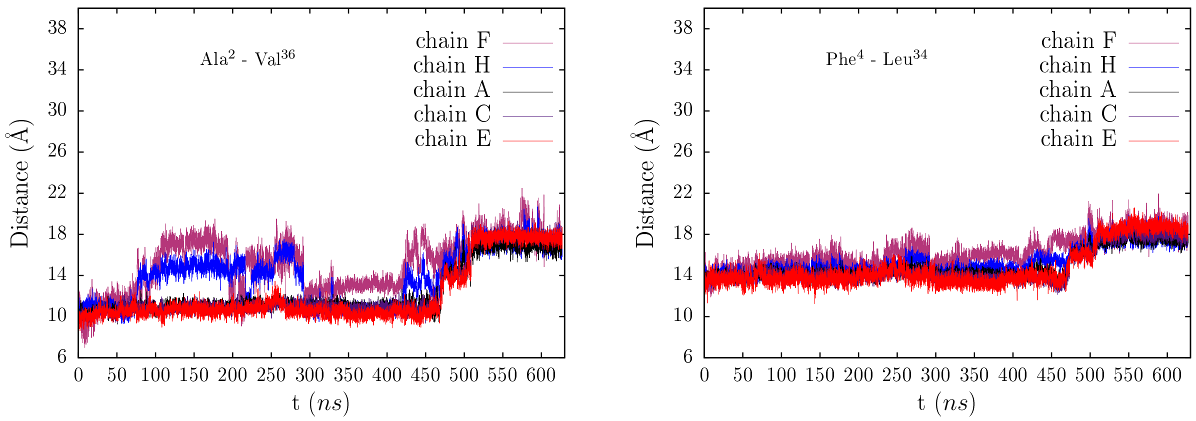


Figure S17: Variation in distance between the residues which form a hydrophobic cluster with time

### Simulation A4, Ligand: $\beta$-aspartyl

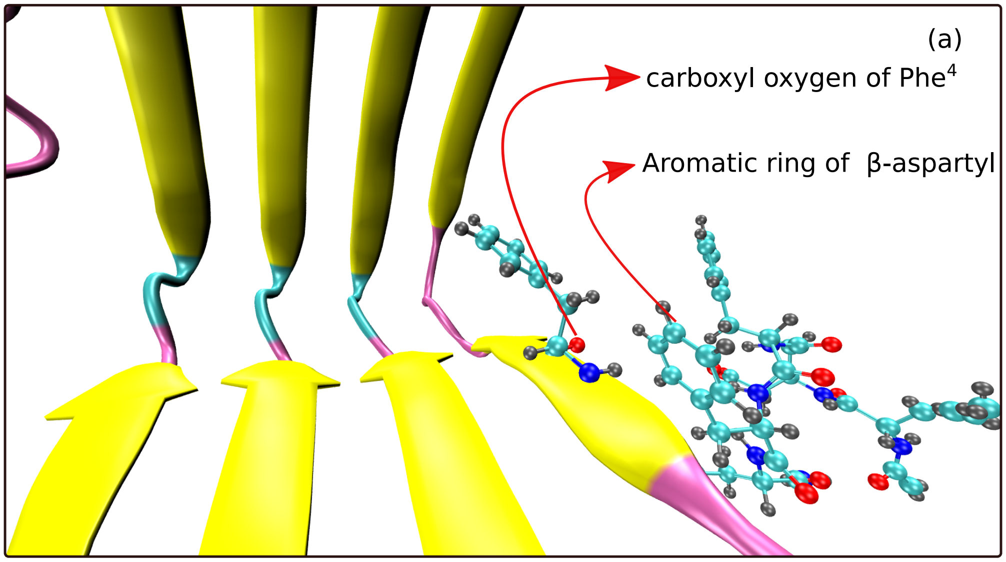


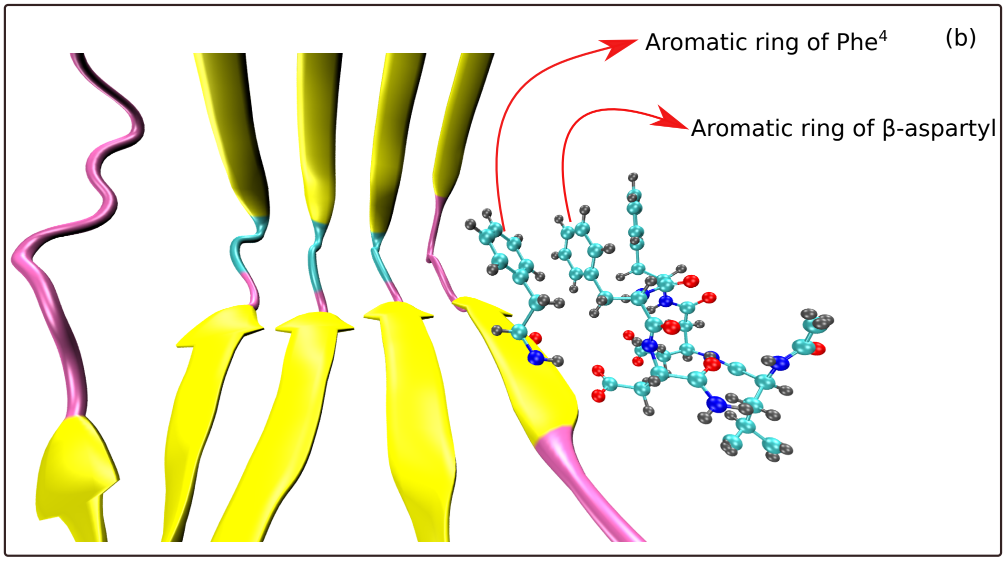


Figure S18: Snapshots showing the sideward allignment which can result in breakage of saltbridges (a) favorable sideward allignment between carboxyl oxygen of $Phe^4$ and aromatic ring of $\beta$-aspartyl, (b) unfavorable sideward allignment between aromatic rings of $Phe^4$ and $\beta$-aspartyl that can lead to the formation of new hydrogen bond between $Asp^1$ and $Gly^{38}$ which further halts the breaking of saltbridges.

### Simulation A4, Ligand: $\beta$-aspartyl

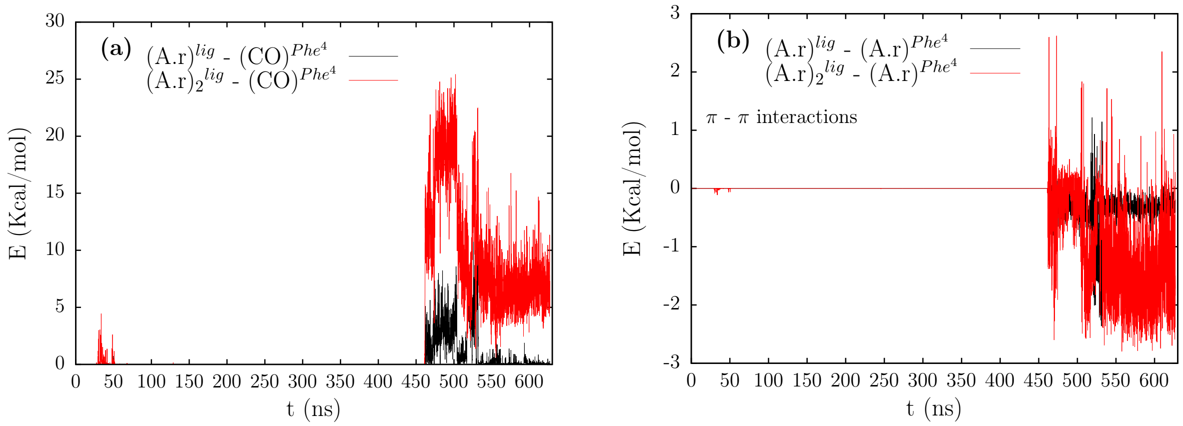


Figure S19: Variation of nonbonded interactions with time between (a) carboxyl oxygen of $Phe^4$ ( $(CO)^{Phe^4}$ ) and two aromatic rings of the ligand, (c) aromatic ring of $Phe^4$ ( $(A.r)^{Phe^4}$ ) and aromatic rings of the ligand

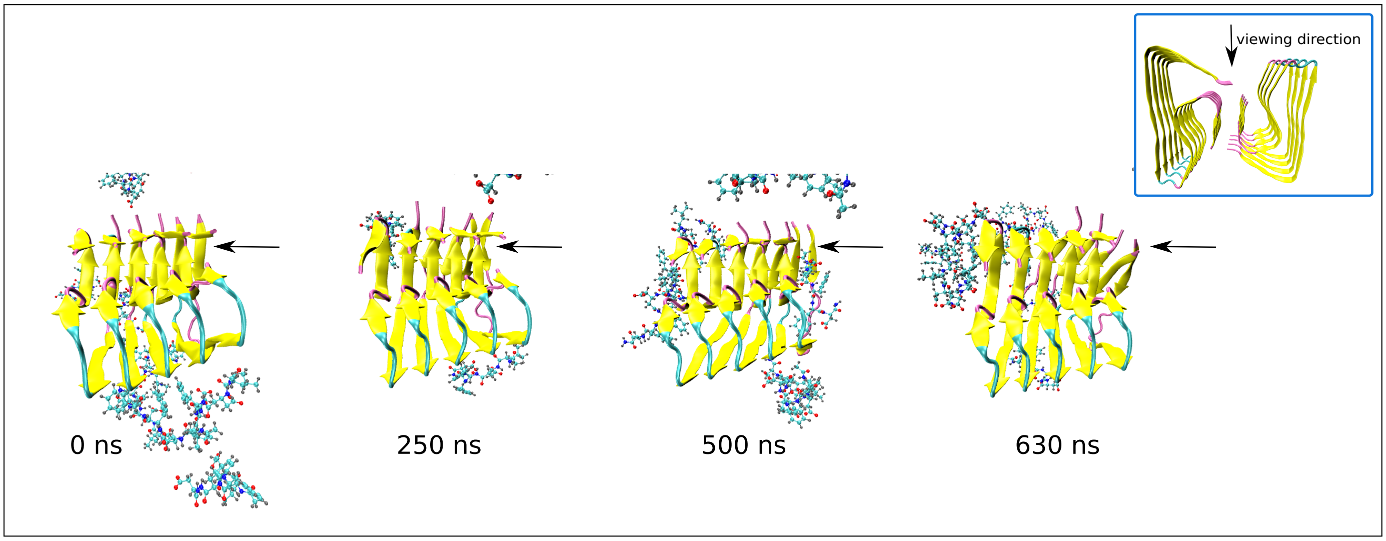


Figure S20: Snapshots showing the topview and the arrow pointing the effect of breakage of saltbridges

### Simulation A5 : Mixture of $\alpha$ & $\beta$ aspartyl in 1:2 ratio

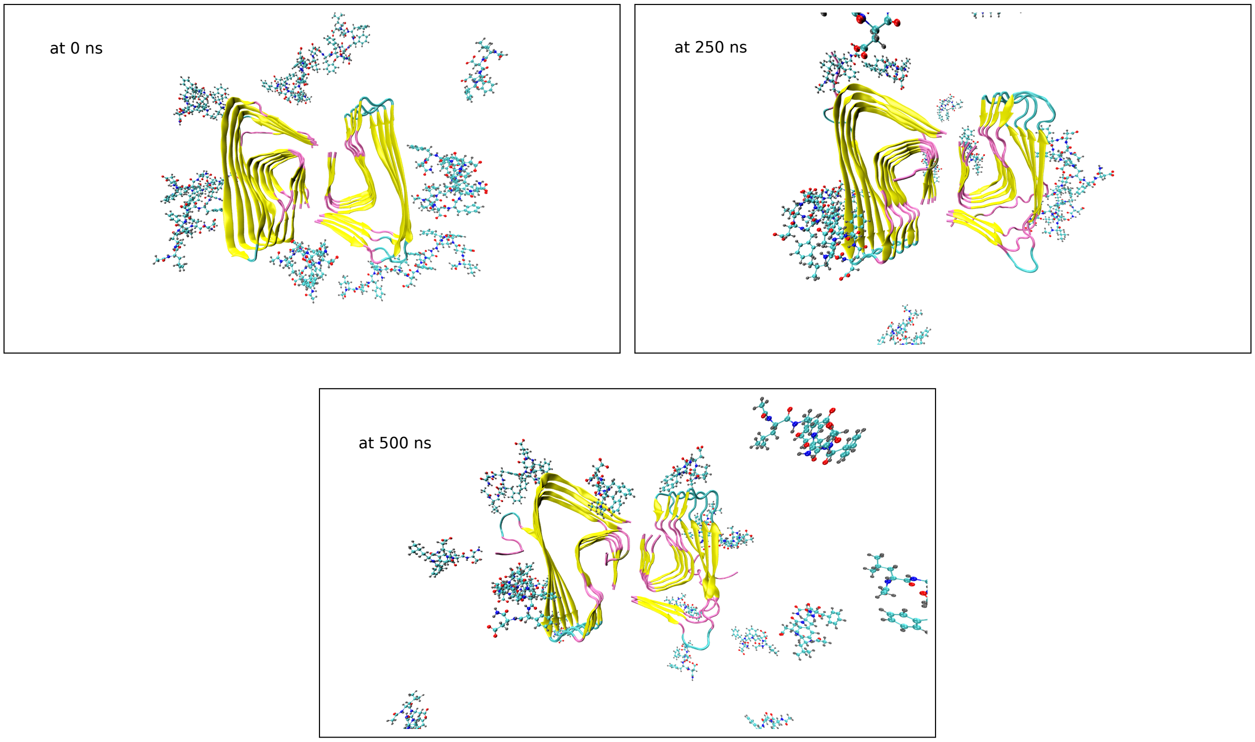


Figure S21: Snapshots showing the protein and the ligands (CPK representation) around it at various times of the simulation

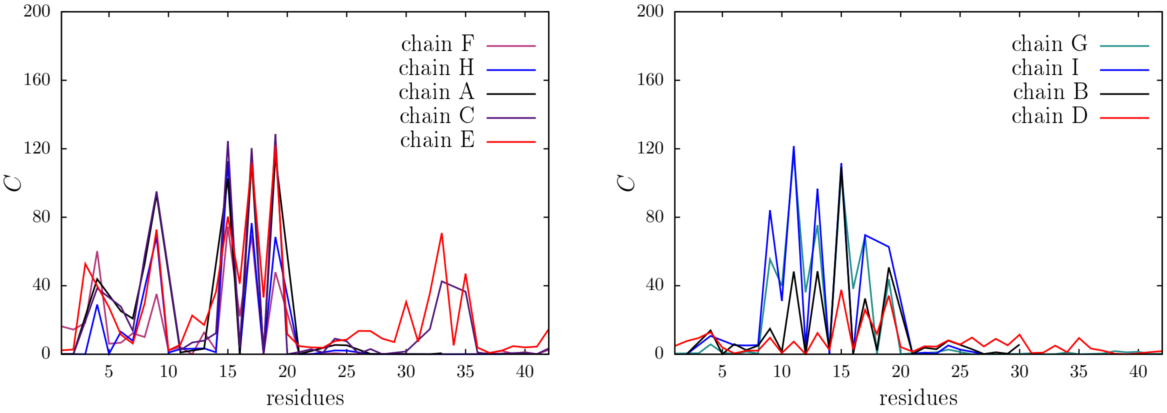


Figure S22: Variation in percentage contacts ($C$) of mixture of $\alpha$ & $\beta$ aspartyl with residues of each subunit of A$\beta$

**Variation in total energy (E) among the ligands in various simulations**

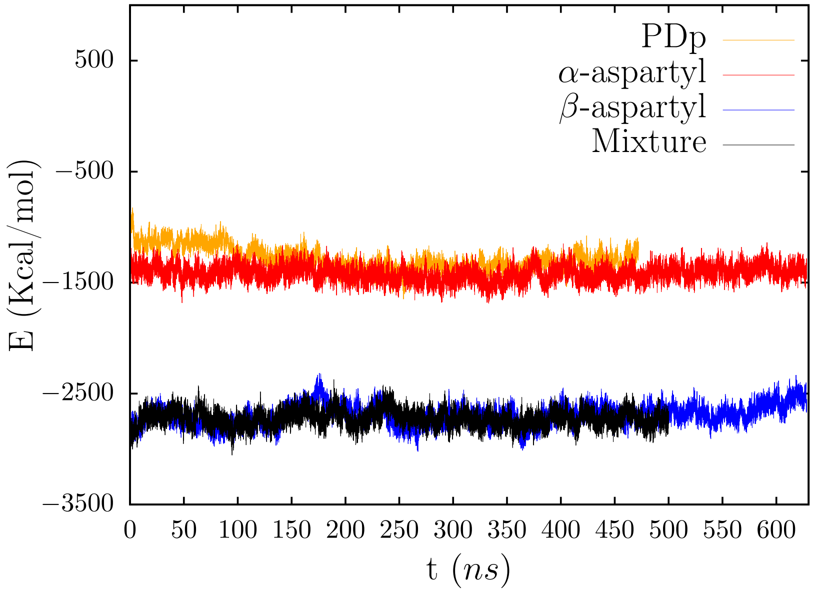


Figure S23
